# Autophagy-dependent gut-to-yolk biomass conversion generates visceral polymorbidity in aging *C. elegans*

**DOI:** 10.1101/234419

**Authors:** Marina Ezcurra, Alexandre Benedetto, Thanet Sornda, Ann F. Gilliat, Catherine Au, Qifeng Zhang, Sophie van Schelt, Alexandra L. Petrache, Yila de la Guardia, Shoshana Bar-Nun, Eleanor Tyler, Michael J. Wakelam, David Gems

## Abstract

Aging (senescence) is characterized by the development of numerous pathologies, some of which limit lifespan. Key to understanding aging is discovery of the mechanisms (etiologies) that cause senescent pathology. In *Caenorhabditis elegans* a major senescent pathology of unknown etiology is atrophy of its principal metabolic organ, the intestine. Here we identify a cause of not only this pathology, but also of yolky lipid accumulation and redistribution (a form of senescent obesity): autophagy-mediated conversion of intestinal biomass into yolk. Inhibiting intestinal autophagy or vitellogenesis rescues both visceral pathologies, and can also extend lifespan. This defines a disease syndrome leading to polymorbidity and contributing to late-life mortality. Activation of gut-to-yolk biomass conversion by insulin/IGF-1 signaling (IIS) promotes reproduction and senescence. This illustrates how major, IIS-promoted senescent pathologies in *C. elegans* can originate not from damage accumulation, but from continued action of a wild-type function (vitellogenesis), consistent with the recently proposed hyperfunction theory of aging.

---

Aging is a pervasive phenomenon across animal species and the leading cause of human morbidity and death worldwide through its associated senescent pathologies. Despite major advances in the genetics of longevity, sparked by the identification of long-lived insulin/IGF-1 signaling (IIS) pathway mutants in the roundworm *Caenorhabditis elegans* (Kenyon 2010), the proximate causes of aging remain unclear. One proposed cause is stochastic molecular damage, due to reactive oxygen species (ROS) for instance (Harman 1956; Beckman and Ames 1998), and consequent accumulation of malfunctioning biomolecules, which IIS promotes (Kenyon 2010; Shore and Ruvkun 2013). Another possible cause of aging is the continued and deleterious action (run-on, or hyperfunction) in later life, of wild-type genes beyond their “intended purpose” (Williams 1957; Blagosklonny 2006; de la Guardia et al. 2016), which we will refer to as the Williams-Blagosklonny theory. This follows from the evolutionary principle of antagonistic pleiotropy (AP), where natural selection can favour gene variants that enhance fitness in early life while promoting senescent pathologies in later life, because the early life benefits to the species outweigh the late-life costs to the individual (Williams 1957). For example, IIS promotes growth and reproduction in early life, and age-related pathologies in late life (Blagosklonny 2010; Gems and Partridge 2013). Conversely, IIS inhibition increases lifespan and healthspan from worms to mammals (Kenyon 2010), but can slow development and reduce fitness.

It now appears that aging-associated molecular damage is more consequence than cause of senescent pathologies (Blagosklonny 2006; Van Raamsdonk and Hekimi 2010; Gems and Partridge 2013), and that multiple aging pathologies can arise from common underlying mechanisms (Gems 2015; Baker et al. 2016). This argues for complementing the standard lifespan genetics approaches that have been so fruitful in invertebrate biogerontology with the study of senescent pathologies, to enhance our understanding of aging. For example, lifespan, traditionally seen as a metric of an underlying continuous aging process (Fig. 1A, top), may alternatively be viewed as a function of one or more life-limiting senescent pathologies. Which of the many extant pathologies limits lifespan may vary between individuals, environmental conditions, genders and species (Fig. 1A, bottom) (Gems 2015; Zhao et al. 2017), confounding studies of the genetics of lifespan. We propose that a pathology-focused approach in aging model organisms may yield new insights into the primary mechanism(s) of aging and help understand how genes control lifespan.

**Figure 1.**
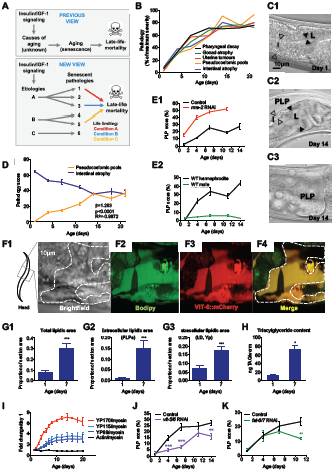
*C. elegans* develops synchronous senescent pathologies from the onset of adulthood, associated with lipoprotein accumulation. (*A*) Top: The traditional view is that lifespan is a metric of an underlying process, aging. Bottom: An alternative view is that lifespan is a function of life-limiting senescent pathologies. (*B*) Senescent pathologies measured by Nomarski microscopy develop in parallel from the onset of adulthood, reaching maximum severity at day 14. (*C*) 14-day old hermaphrodites (*C2-3*) exhibit an atrophied intestine (hollow arrowheads), an enlarged lumen (L, black arrowheads), and more pseudocoelomic pools compared to 1-day old worms (*C1*). (*D*) The evolution of PLP accumulation mirrors intestinal atrophy, and both are strongly correlated at a population level (β: regression slope, *p*: p-value, R: correlation coefficient). (*E*) Inhibition of yolk uptake by *rme-2* RNAi aggravates PLP accumulation (*E1*), while absence of yolk synthesis prevents PLP accumulation in aging males (*E2*), n=2 trials. (*F*) PLPs contain neutral lipids and yolk proteins YP115/YP88 as revealed by co-localization (*F4*) of Bodipy (*E2*) and VIT-6::mCherry (*F3*). (*G*) Estimation of lipid content from measuring intracellular yolk particle (Yp), lipid droplet (LD), and extracellular PLP areas in TEM of mid-body sections reveals a 3.75-fold increase between day 1 and 7 (*G1*), mostly explained by a 15-fold rise in combined PLP area (*G2*). (*H*) Colorimetric assays for triglyceride detection show an 8-fold increase in TAG content between day 1 and 7 of adulthood. (*I*) Analysis of Coomassie stained SDS-PAGE of wildtype worm lysates (Depina et al. 2011) shows that YP170, YP115 and YP88 content dramatically increases with age. (*J*) Inhibition of YP synthesis by *vit-5/-6* RNAi reduces age-associated PLP accumulation. (*K*) Inhibition of yolk lipid synthesis by *fat-6/-7* RNAi lowers PLP score. Experiments were performed at 20°C without FUDR. Worms were raised on *E. coli* B OP50, except for panels (*D*), (*I*) and (*J*), where *E. coli* K12 HT115 was used. * *p*<0.05, ** *p*<0.01, *** *p*<0.001.

*C. elegans* exhibits many prominent senescent pathologies, including degeneration of the pharynx, tumors in the uterus, atrophy of the intestine and gonad, and steatotic lipoprotein redistribution (Garigan et al. 2002; Herndon et al. 2002; Luo et al. 2010; Hughes et al. 2011; McGee et al. 2011; de la Guardia et al. 2016; Palikaras et al. 2017), thus providing ample material for study. Here we have focused on elucidating senescent pathologies of the main *C. elegans* metabolic organ, the intestine, which fulfills the functions of liver and adipose tissue and is a main site of action for IIS-mediated effects on lifespan (Libina et al. 2003; Budovskaya et al. 2008). To this end we employed a *developmental pathology* approach, quantifying the temporal evolution of gut pathologies at anatomical, cellular and molecular levels, and identified a novel pathophysiological mechanism: autophagy-dependent gut-to-yolk biomass conversion, run-on of which causes multiple visceral pathologies, including intestinal atrophy. These results are consistent with the account provided by the Williams-Blagosklonny view of how antagonistic pleiotropy is enacted in terms of proximate mechanisms.

## Results

### Major senescent pathologies develop in early to mid-adulthood and limit life

Various senescent pathologies have been documented in *C. elegans*, but details of their developmental timing, particularly in relation to one another, remain sparse. In humans, diseases of aging (e.g. cardiovascular, neurodegenerative, cancer) increase mainly towards the end of life. We asked, is this true of *C. elegans* too? To this end, we monitored development of uterine tumors, gonadal atrophy, deterioration of the pharynx (foregut), yolky lipoprotein pools in the body cavity (pseudocoelom), and intestinal atrophy. Wild-type (N2) hermaphrodites were aged under standard conditions (NGM agar plates, *Escherichia coli* OP50 bacteria as a food source, 20°C), imaged using Nomarski microscopy at intervals from day 1 to 21 (d1-21) of adulthood, and the severity of pathologies was quantified (Supplemental Fig. S1A-E). Against expectation, senescent pathologies developed not towards the end of life but much earlier, with initiation at around the time of self-sperm depletion (d3-4) and peak severity reached before the median lifespan (Supplemental Fig. 2A). Moreover, the various pathologies developed in relative synchrony, revealing an aging syndrome (Fig. 1B). In particular, development of intestinal atrophy and pseudocoelomic lipoprotein pool (PLP) accumulation (Fig. 1C) was strongly coupled in all growth conditions tested, showing correlation within individuals (Supplemental Fig. 2B, C) and strikingly tight temporal correlation (Fig. 1D). This suggested a possible common etiology of these two pathologies, which we investigated further, beginning with a closer examination of each pathology individually.

### Accumulation of pseudocoelomic lipoprotein pools as a form of senescent obesity in C. elegans

PLPs have been previously identified as yolk as they contain the yolk protein (vitellogenin) VIT-2/YP170 (Garigan et al. 2002; Herndon et al. 2002). We verified this by co-localization within the pools of both VIT-2 and VIT-6 (the sole source of YP115 and YP88 in worms, Supplemental Fig. 2D). Consistent with PLPs being yolk, preventing oocyte yolk uptake by RNAi against the receptor *rme-2* (Grant and Hirsh 1999) accelerated pool growth, while males (which do not make yolk) did not accumulate PLPs (Fig. 1E). The magnitude of the pools, which can grow to fill the entire body cavity (Fig. 1C3), and the lipid content of yolk (Sharrock et al. 1990), implies a major senescent buildup of lipids in *C. elegans*. Fluorescent labeling of neutral lipids using gentle-fix Bodipy staining (Klapper et al. 2011) confirmed the presence of lipids in PLPs, which co-localized with VIT-6::mCherry (Fig. 1F, Supplemental Fig. 2E). Moreover, transmission electron microscopy (TEM) of mid-body sections defined a 3.8-fold increase in lipid organelle area between day 1 and 7 of adulthood (Fig. 1G1), largely attributable to a 15-fold increase in pool area (Fig. 1G2), but also to a 3-fold increase in intracellular lipids (Fig. 1G3). During the same period a striking eight-fold increase in triglyceride (TAG) content was detected, using biochemical assays and lipidomic analysis (Fig. 1H, Supplemental Fig 2F-G1, Supplemental Table 2). Thus, aging *C. elegans*, by accumulating large amounts of ectopic fat within the pools, become steatotic (Palikaras et al. 2017).

### Run-on of yolk synthesis promotes yolky pool generation

PLP accumulation has been suggested to result from run-on of lipoprotein production after cessation of egg laying (continued source but loss of sink) (Herndon et al. 2002). To characterize this putative vitellogenic open faucet, we monitored YP levels throughout life (days 1-20). This revealed a sustained increase in YP content until day 14, and a maximal 7-fold increase in YP170 (Fig. 1h, Supplemental Fig. 3C). Blocking YP accumulation by *vit-5/-6* double RNAi (Supplemental Fig. 3D) or inhibiting yolk lipid synthesis by RNAi against the fatty acid desaturases *fat-6* and *fat-7* (Watts and Browse 2002; Li et al. 2011) reduced pool accumulation (Fig. 1J-K), confirming that PLPs stem from run-on of lipoprotein synthesis.

### Conversion of intestinal biomass into yolk causes organ atrophy

While cessation of egg laying after sperm depletion is a pre-requisite for YP accumulation (Depina et al. 2011) post-reproductive accumulation must reflect continued vitellogenin synthesis. It is notable that *C. elegans* is able to sustain heavy production of lipoprotein to advanced ages, despite the decline with age in feeding rate (Huang et al. 2004) (Supplemental Fig. 3A). We also noted that while *C. elegans* total protein content plateaus at day 5, the amount of vitellogenin as a proportion of overall protein continues to increase for several more days; strikingly, vitellogenin eventually forms 30-40% of total worm protein content (Fig. 2A-B, Supplemental Fig. 3B). These observations, taken together with the coupling between intestinal atrophy and pool accumulation (Fig. 1B-D) in all growth conditions tested (altered temperature or bacterial diet, presence of FUDR, Supplemental Fig. 2B-C), and the fact that the intestine is the site of yolk production (Kimble 1983) suggest a new hypothesis: that the intestine consumes its own biomass to enhance capacity for yolk production.

**Figure 2.**
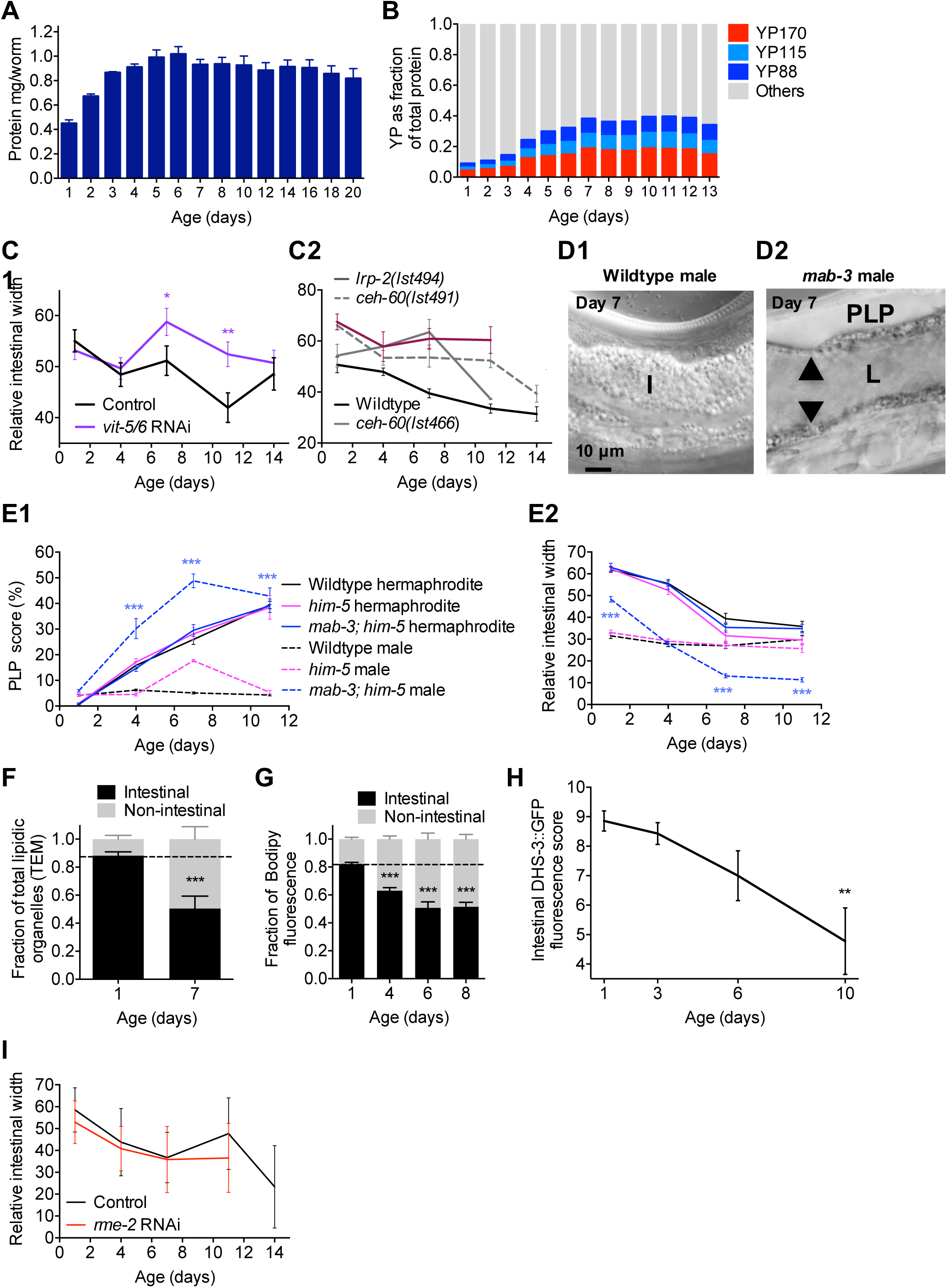
Run-on of yolk production in aging worms entrains intestinal atrophy via conversion of intestinal mass into lipoprotein. (*A*) Total protein content peaks at day 5, doubling from day 1, in wildtype animals. (*B*) The proportion of YP to total proteins measured by densitometry analysis increases 4-folds between days 1 and 7. (*C*) Inhibition of yolk synthesis by *vit-5/-6* RNAi (*C1*), *lrp-2* or *ceh-60* mutations (*C2*) rescues intestinal atrophy, *n*=2 trials. All mutants statistically significant different from wildtype at least two time points, p<0.05. (*D*) Wildtype males are devoid of gut pathologies (*D1*) while yolk-producing *mab-3* (*Doublesex/DMRT1* homologue) males display PLPs and gut atrophy. (E) Ectopic yolk production in *mab-3* males induces ectopic and very severe age-associated PLP accumulation (E1) and gut atrophy (E2), n=2 trials. Experiments performed in the male generating background *him-5*. Statistical significance between *mab-3; him-5* and *him-5* males. (*F*) TEM of mid-body sections reveals a redistribution of intestinal lipids to other tissues between day 1 and 7, confirmed by analysis of neutral lipids staining by Bodipy (*G*), and consistent with a decrease in intestinal lipid droplets revealed by DHS-3::GFP labeling in older age (*H*). (*I*) Inhibition of yolk uptake by *rme-2* RNAi, which aggravates PLP accumulation, does not affect intestinal atrophy. Experiments conducted at 20°C without FUDR (*A-F, I*) and 25°C with FUDR (*G, H*). * *p*<0.05, ** *p*<0.01, *** *p*<0.001.

Consistent with this hypothesis, inhibition of YP production by *vit-5/-6* RNAi (Supplemental Fig. 3D) or by mutation of upstream activators of yolk synthesis (*ceh-60* and *lrp-2*) (Rompay et al. 2015) rescued both pool accumulation and intestinal atrophy (Fig. 1J, 2C, Supplemental Fig. 3E). Moreover, in wild-type males, which do not make yolk, neither PLP accumulation nor intestinal atrophy were observed (Fig. 2D-E). However, induction of ectopic YP production in males by mutation of *mab-3* was sufficient to induce both pathologies (Fig. 2D-E). Corroborating the idea that intestinal mass is converted into lipoprotein, analysis of lipid distribution by TEM and Bodipy staining revealed that the proportion of intestinal lipids vs PLP lipids decreases between day 1 and 7 (Fig. 2F-G), while DHS-3::GFP labeling of intestinal lipid organelles (Na et al. 2015) becomes patchy and reduced (Fig. 2H, Supplemental Fig. 3F-G). Consistent with this, inhibition of YP production by *vit-2* RNAi increases intestinal lipid content (Seah et al. 2016). An alternative possibility is that steatosis rather than yolk synthesis indirectly causes intestinal damage and atrophy. *rme-2* RNAi, which increases steatosis by inhibiting yolk uptake by oocytes, did not aggravate intestinal atrophy (Fig. 2I) thus arguing against this; a further possibility is that intestinal activation of the SKN-1/Nrf2 transcription factor by *rme-2* RNAi (Steinbaugh et al. 2015) impedes atrophy.

Altogether, these results imply that a single etiology, intestinal biomass conversion into lipoprotein, causes three comorbidities in senescing *C. elegans*: intestinal atrophy, extracellular yolk accumulation and steatotic lipid redistribution.

### Autophagy promotes intestinal atrophy and PLP accumulation by gut biomass conversion

But how does this biomass conversion occur? Such a process would require bulk breakdown and recycling of cellular components, a role typically fulfilled by autophagy (macroautophagy, specifically) in contexts of starvation, ecdysozoan molting and metamorphosis (Cuervo 2004). Notably, levels of autophagy appear to be high in the intestine of adult hermaphrodites (Chapin et al. 2015). To test this hypothesis we first examined mutants defective in induction of autophagy (*atg-13(bp414)*), autophagosomal vesicle elongation (*atg-4.1(bp501)*), and Atg9p retrieval (*atg-2(bp576)* and *atg-18(gk378)*), and found that all four mutations reduced both gut atrophy and pool accumulation (Fig. 3A). This implies that autophagy promotes both pathologies.

**Figure 3.**
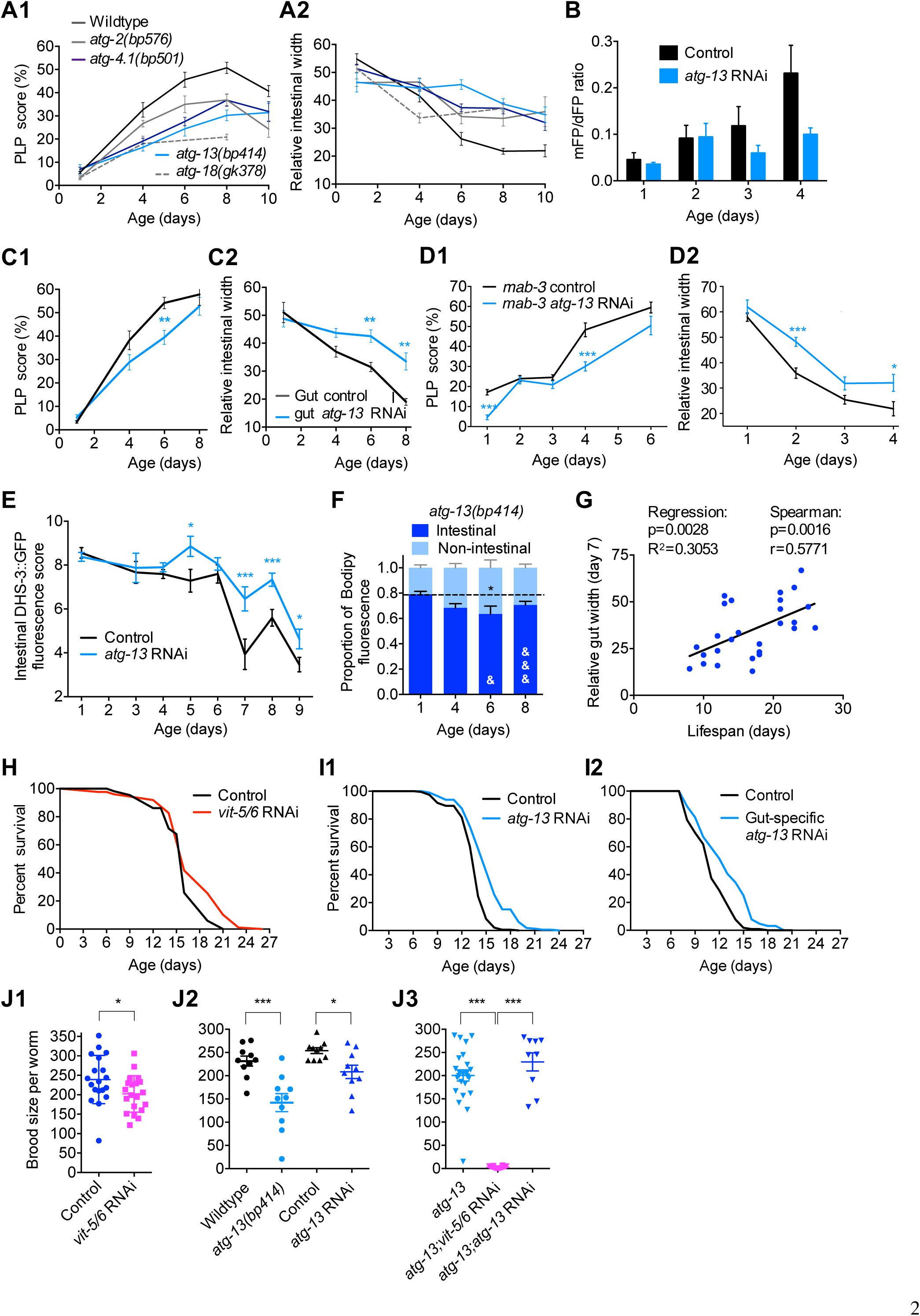
Intestinal autophagy mediates gut-to-yolk biomass conversion, promoting optimal reproduction at the expense of longevity. (*A*) Mutations in autophagy genes reduce age-associated PLP accumulation (A1) and limit intestinal atrophy (A2), *n*=2 trials. For all mutants, at least two time points difference was statistically significant, p<0.05. (*B*) *atg-13* RNAi suppresses the age increase in intestinal autophagy (*p*=0.009, two-way ANOVA). (*C*) Adult-limited *atg-13* RNAi targeted to the gut reduces age-associated PLP accumulation (*C1*), and intestinal atrophy (*C2*), *n*=2 trials. (*D*) Adult-limited *atg-13* RNAi rescues age-associated ectopic PLP accumulation and gut atrophy in *mab-3* males. (*E*) Adult-limited *atg-13* RNAi delays the age-associated decrease in DHS-3::GFP-labelled intestinal lipid droplets, and (F) inhibits age-associated lipid redistribution. & *p*<0.05, &&&, *p*<0.001, compared to age-matched wildtype (Fig. 2G). (*G*) Individual worm pathology analysis indicates that intestinal width at day 7 correlates positively with longevity. (*H*) Inhibition of yolk synthesis by *vit-5/-6* RNAi extends lifespan, *n*=5 trials. (*I*) Adult-limited *atg-* 13 RNAi targeted to the whole soma (*I1*) n=6 trials, or to the gut only (*I2*), *n*=5 trials, can extend lifespan. (*J*) Adult-limited *vit-5/-6* RNAi (*J1*) or *atg-13* RNAi, and *atg-13* mutation (*J2*) each modestly reduce brood size, but *vit-5/-6* RNAi and *atg-13* mutant synergistically cause sterility (*J3*), *n*=2 trials. * *p*<0.05, ** *p*<0.01, *** *p*<0.001. experiments performed at 20°C without FUDR (*G, H*) and 25°C with FUDR (*A-F, I, J*). Note that the *atg-2* mutation is in the *him-5* background.

To explore this further we tested effects of adult-specific inhibition of autophagy by *atg-13* RNAi, selecting *atg-13* (Tian et al. 2009) since as an upstream regulator of autophagy its inhibition may be less likely to induce deleterious pleiotropic effects through accumulation of abnormal autophagosomes. Using an intestine-specific reporter of autolysosome formation (Chapin et al. 2015), we first obtained evidence that adult-specific *atg-13* RNAi reduces intestinal autophagy (Fig. 3B). *atg-13* RNAi also reduced yolk pool accumulation, and intestinal atrophy (Supplemental Fig. 4A). Notably, *atg-13* RNAi restricted to the adult intestine also suppressed both pathologies (Fig. 3C), while gut-specific transgenic rescue of the *atg-13(bp414)* mutation restored intestinal atrophy (Supplemental Fig. 4B). Moreover, adult-limited *atg-13* RNAi rescued pathologies induced by intestinal feminization in *mab-3* males (Fig. 3D).

Abrogation of *atg-13, atg-4.1, atg-9, atg-2* or *atg-18* by means of mutation or RNAi also impeded senescent lipid redistribution, as shown by DHS-3::GFP, Bodipy and TEM lipid organelle analyses (Fig. 3E-F, Supplemental Fig. 4C-H). Moreover, *atg-13(bp414)* largely suppressed age changes in lipid profiles (Supplemental Fig 4K, Supplemental Table 2). As autophagosomes consume endomembranes, we followed gut intracellular markers for Golgi, early, late and recycling endosomes during early aging. We failed to see salient age-related changes in endosomal staining, but we did observe an age-dependent reduction in intestinal Golgi labeling by alpha-mannosidase II-GFP (Rolls et al. 2002), which was rescued by *atg-13* RNAi (Supplemental Fig. 4J). This would be consistent with the hypothesis that autophagosomes compete with Golgi apparatus for endomembrane availability (Ge et al. 2013). These results suggest that atg-13-mediated lipophagy promotes the conversion of intestinal lipids into yolk lipids destined for export. A caveat here is that early inhibition of other autophagy genes during development has been shown to reduce lipid levels in young adults (Lapierre et al. 2013), implying that autophagy can also promote lipid storage by unknown pathways. However, given that blocking autophagy inhibits rather than increases loss of adult intestinal biomass, a parsimonious explanation is that such effects are attributable to the canonical, catabolic action of autophagy. Taken together, our results suggest that autophagy promotes gut-to-yolk biomass conversion, leading to senescent polymorbidity.

### Gut biomass conversion involves a trade-off between fitness and late-life health

We then explored whether gut-to-yolk biomass conversion contributes to late life mortality. Analysis of age-specific pathology measures and survival in individual worms identified a negative correlation between lifespan and intestinal atrophy in mid-life (Fig. 3G), suggesting that intestinal atrophy contributes to mortality. Consistent with this, inhibiting gut-to-yolk biomass conversion by *atg-13* RNAi targeted to the whole adult animal or to the intestine alone consistently extended lifespan at 25°C (+21.3% and +5.4%, respectively, Fig. 3I, Supplemental Fig. 4L, Table 1), though not at 20°C (Supplemental Table 1). Lifespan was also extended by adult-limited, whole worm RNAi of *atg-2*, but not *atg-18* (25°C, Supplemental Fig. 4L, Table 1). One possibility is that this reflects condition-dependency with respect to which senescent pathologies are life-limiting (Fig. 1A). *vit-5/-6* RNAi also consistently increased lifespan (+12.8%, Supplemental Table 1), as reported for *vit-5* RNAi (Murphy et al. 2003).

Although *vit-5/-6* RNAi, *atg-13* RNAi and *atg-13(bp414)* had little effect on brood size, consistent with the normal fertility of *ceh-60* and *lrp-2* mutants (Rompay et al. 2015), *vit-5/-6* RNAi in *atg-13(bp414)* worms caused near-sterility (Fig. 3J). These results suggest that high levels of autophagy in the intestine promote fitness by supporting yolk production in early adulthood (during reproduction), while non-adaptive run-on of intestinal autophagy contributes to gut atrophy and pool formation in later life (post-reproduction). This is further supported by the fact that mated hermaphrodites with increased brood sizes display accelerated gut atrophy (Supplemental Fig. 5A) consistent with life-shortening effects of exposure to males (Shi et al. 2017), but exhibit delayed PLP accumulation, likely due to prolonged egg laying (increased yolk sink, Supplemental Fig. 5B). Exposure to male secretions alone did not affect either visceral pathology (Supplemental Fig. 5C-H), consistent with recent evidence that exposure to males shortens lifespan by two distinct mechanisms (Shi et al. 2017). All this supports the existence of a trade-off between beneficial effects of biomass conversion on reproductive fitness, and later pathogenic effects.

### Insulin/IGF-1 signaling (IIS) promotes gut-to-yolk biomass conversion

Another factor that enhances reproductive fitness at the expense of senescent pathologies and lifespan shortening is IIS (Garigan et al. 2002; Kenyon 2010). Hence, the wild-type *daf-2* insulin/IGF-1 receptor allele is a classic example of AP (Blagosklonny 2010). The proximate mechanisms downstream of IIS that account for effects on lifespan have been sought for many years, e.g. by detailed characterization of IIS regulated genes, yet remain unclear. One possibility is that gut-to-yolk biomass conversion is one of these proximate mechanisms. Consistent with this idea, mutation of *daf-2* not only extends lifespan (Kenyon 2010) but also rescues senescent polymorbidity, including YP increase, PLP accumulation, intestinal atrophy, and lipid redistribution (Fig. 4A-D, Supplemental Fig. 6A-D) (Garigan et al. 2002; Depina et al. 2011). This rescue requires the DAF-16/FoxO transcription factor (Fig. 4C-D. Supplemental Fig. 6D) (Kenyon 2010; Depina et al. 2011). Moreover, *atg-13* RNAi extends the short lifespan of *daf-16(mgDf50)* (Fig. 4E), suggesting that autophagy acts downstream of or in parallel to DAF-16. Understanding how IIS regulates gut-to-yolk biomass conversion will require further study.

**Figure 4.**
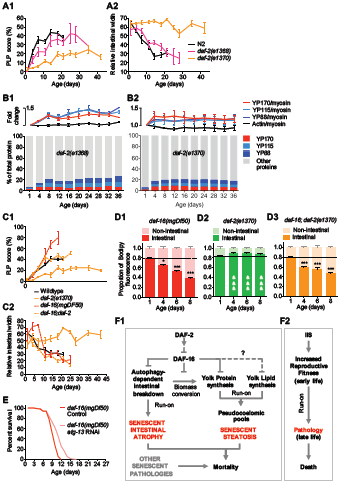
Reduced insulin/IGF-1 signaling inhibits gut-to-yolk biomass conversion. (*A*) PLP accumulation (*A1*) and intestinal atrophy (*A2*) are both suppressed in *daf-2(e1370)* and *daf-2(e1368)* mutants. (*B*) YP levels and YP proportion in *daf-2(e1368)* (*B1*) and *daf-2(e1370)* (*B2*). (*C*) PLP accumulation (*C1*) and intestinal atrophy (*C2*) in *daf-2(e1370)* require *daf-16*, *n*=2 trials. (*D*) Age-associated neutral lipid redistribution revealed by Bodipy fluorescence is suppressed in *daf-2(e1370)* mutants and requires *daf-16*. * *p*<0.05, *** *p*<0.001. &&& *p*<0.001, compared to age-matched *daf-16, daf-16; daf-2*, or wildtype (Fig. 2G). (F) Proposed mechanism of antagonistic pleiotropy action of IIS promoting reproduction at the expense of senescent pathologies, through autophagy-mediated gut-to-yolk biomass conversion.

## Discussion

These findings support a model in which the run-on of reproductive functions promotes senescent pathogenesis in *C. elegans* hermaphrodites. This involves IIS-driven and autophagy-mediated conversion of intestinal biomass into steatotic yolk pools. The model proposes a single etiology for major comorbidities of worm aging, and identifies a mechanism by which IIS, through inhibition of DAF-16, can cause senescent atrophy of the hermaphrodite intestine (Fig. 4F, Supplemental Fig. 6E).

The mechanisms proposed here are at odds with the traditional view of aging as caused by homeostatic imbalance and damage accumulation. Here instead, persistence of wild-type gene function beyond their “intended” purpose actively drives senescent pathogenesis, consistent with the hyperfunction model of AP action (Williams 1957; Blagosklonny 2006). This contrasts with the more traditional, damage-based, disposable soma model of AP action, which postulates that genes can promote fitness by diverting resources from somatic maintenance to reproduction, thereby accelerating damage accumulation and, consequently, senescence (Kirkwood and Rose 1991).

### Coordinated development of senescent pathologies in mid-adulthood

A survey of patterns of development of major senescent pathologies revealed that these appeared not, as expected, near the end of life, but in mid-adulthood. The overall pattern suggests a mid-life phase characterized by vigorous development of pathologies, ending around d10-12 of adulthood (Fig. 1B). The major anatomical changes that result likely contribute to reported age-changes in *C. elegans* expression profiles, e.g. levels of intestinal proteins decline with age (Walther et al. 2015). Thus, many age changes in mRNA and protein level may be the consequence rather than the cause of pathogenic processes such as gut-to-yolk biomass conversion. One possibility is that when the major pathologies have reached their developmental endpoints around d10-12 the nematodes are, as a consequence, terminally ill and spend the rest of their lives slowly dying (Podshivalova et al. 2017).

### Yolky pools as a form of senescent obesity in C. elegans

Yolk accumulation in adult *C. elegans* has been described previously, but not its full extent during aging. Here we report that age increases in vitellogenin levels reach 7-fold that of young adults, forming up to 30-40% of total worm protein, while TAG levels increase ~8 fold. The presence of large yolky pools staining with the lipid dye Bodipy implies that such pools are major extracellular lipid deposits, helping to account for the age increase in TAG. The more modest age-associated increases in other lipids found in yolk might reflect concomitant reductions in their levels due to organ atrophy. The presence in elderly nematodes of large, extracellular lipid pools could have been overlooked previously due to the use in lipid staining protocols of fixation methods with dehydration steps; here we used a gentle staining methods without such steps (Klapper et al. 2011).

We postulate that yolky pools develop by run-on of yolk lipid production (Fig. 4F), but cannot exclude a contribution from lipids released by degeneration of other organs such as the distal gonad (de la Guardia et al. 2016). The age-dependent accumulation and gross appearance of the yolky pools suggest that they represent a form of pathological senescent obesity in *C. elegans.* Consistent with this, lifespan was reduced by knock-down of expression of the yolk receptor *rme-2*, which accelerated yolky pool accumulation without affecting intestinal atrophy (Fig. 1E and 2I). A possible explanation for yolky pool toxicity is late-life ectopic deposition e.g. in muscle and epidermis (data not shown), due to yolky pool endocytosis as part of a senescent redistribution of fat stores. This age-associated redistribution of fat is reminiscent of pathogenic shifts in fat distribution that occur in aging mammals (Huffman and Barzilai 2010), underscoring the potential of *C. elegans* as a model for investigating pathologies of fat metabolism (Jones and Ashrafi 2009).

### Gut-to-yolk biomass conversion causes life-limiting polymorbidity

The intestine is the major somatic internal organ in *C. elegans*, also serving as liver and adipose tissue. As *C. elegans* hermaphrodites get old, the intestine undergoes severe deterioration (Haithcock et al. 2005; McGee et al. 2011), likely impairing worm viability. Consistent with this, the extent of intestinal atrophy in early-mid adulthood is predictive of lifespan (Fig. 3G). Furthermore, the lifespan-controlling transcription factors DAF-16, SKN-1 and ETS-4 exert their effects in the intestine (Libina et al. 2003; Tullet et al. 2008; Thyagarajan et al. 2010). Moreover, intestinal necrosis is a key event in organismal death (Coburn et al. 2013). All this is consistent with the view that intestinal senescent pathology can be life-limiting to *C. elegans*. Our study defines a likely major cause of intestinal senescence: gut-to-yolk biomass conversion.

In mammals, senescence causes diverse pathologies. Important questions are: to what extent do multiple senescent pathologies originate from common etiologies (Gems 2015)? And: what is the nature of these etiologies? Here we provide an example of how one etiology, gut-to-yolk biomass conversion, promotes two major senescent pathologies: intestinal atrophy and yolk accumulation, each of which may lead to further pathologies. For example, yolk accumulation promotes age-associated vulval integrity defects (Leiser et al. 2016) and uterine tumor development (Wang et al. 2017).

### Autophagy can promote senescent visceral pathologies

Autophagy is traditionally viewed as protective against aging, consistent with the molecular damage theory, and life-shortening effects of inhibition of autophagy (Gelino and Hansen 2012). However, autophagy can both inhibit and enhance the development of pathologies (Shintani and Klionsky 2004). Here we describe a seemingly autophagy-dependent process (gut-to-yolk biomass conversion) that promotes several senescent pathologies in *C. elegans* hermaphrodites, which can be life-limiting. Notably, adult-limited inhibition of *atg-13* consistently extended lifespan at 25°C, though not at 20°C. We postulate that the pathologies that limit life vary with genotype and culture conditions. Moreover the effect on lifespan of inhibiting autophagy may depend on whether life-limiting pathologies are inhibited or enhanced by autophagy, and on severity or timing of inhibition. For example, it was recently shown that late-life inhibition of neuronal autophagy can increase *C. elegans* lifespan (Wilhelm et al. 2017) (see Supplemental Discussion).

### *Gut-to-yolk biomass conversion: a mechanism for* daf-2 *antagonistic pleiotropy*

This study suggests a mechanism by which IIS causes visceral pathology: autophagy-dependent gut-to-yolk biomass conversion (Fig. 4F1). This is different from a prior suggestion that inhibition of autophagy by IIS promotes aging (see Supplemental Discussion).

Aging evolves as the result of antagonistic pleiotropy (AP), implying that late-life gene action causes senescence, including senescent pathologies (Williams 1957; Blagosklonny 2006). *daf-2* is a gene with strong AP effects, promoting reproductive growth in early life, and senescent pathology in later life (Blagosklonny 2010; Gems and de la Guardia 2013). The insight that IIS promotes gut-to-yolk biomass conversion provides an account of a mechanism by which IIS-determined AP is enacted, as follows. In early adulthood, IIS promotes gut-to-yolk biomass conversion, which increases reproductive fitness by increasing yolk production capacity. Supporting this, combined knockdown of autophagy and vitellogenesis strongly reduces fertility (Fig. 3J3) and protein turnover rate is reduced in *daf-2* mutants (Depuydt et al. 2013; Stout et al. 2013; Dhondt et al. 2016). Continued gut-to-yolk biomass conversion later in adulthood leads to pathologies in the form of intestinal atrophy and yolky pools, which can shorten lifespan (Fig. 4F2).

An interesting question is: why do hermaphrodites not turn off gut-to-yolk biomass conversion after cessation of reproduction? Here the evolutionary theory suggests an answer: that such an off-switch did not evolve due to the reduced force of natural selection in post-reproductive animals. By this view, biomass conversion is simply left on, in an open faucet mechanism, an example of process run-on or “hyperfunction” (Herndon et al. 2002; Blagosklonny 2006; de la Guardia et al. 2016), where continued yolk production after reproduction in self-fertilizing hermaphrodites is futile in fitness terms. However, it remains possible that later yolk production does promote fitness e.g. in mated hermaphrodites, which have a longer reproductive period.

Run-on also contributes to other *C. elegans* senescent pathologies that are promoted by IIS, including run-on of embryogenetic programs in unfertilized oocytes which promotes tumor development (McGee et al. 2012; Wang et al. 2017), and germline apoptosis which promotes gonad atrophy (de la Guardia et al. 2016). That IIS promotes mechanistically very distinct hyperfunctions as part of the hermaphroditic reproductive program suggest that run-on mechanisms may be typical of IIS. Given the conserved role and AP effects of IIS in animal aging, one could expect similar mechanisms to contribute to human aging (Rodriguez et al. 2017).

### Conclusions

This study defines a syndrome of senescent polymorbidity driven by wild-type gene action in which atrophy of one tissue is coupled by ectopic fat deposition to pathology in other tissues. Possible generalization of this type of etiology is suggested by parallels with bone erosion in lactating mammals, which ensures sufficient calcium in milk (Hopkinson et al. 2000). In humans, menopause-associated run-on of such bone-to-milk calcium transfer may contribute to both osteoporosis and conditions promoted by ectopic Ca^2+^ deposition, such as vascular calcification and osteoarthritis. More broadly, these investigations demonstrate how senescent pathologies can be generated by the futile sustained activity of naturally selected genetic programs (Williams 1957; Blagosklonny 2006), rather than by passive and stochastic wear and tear processes.

## Materials and methods

### Culture methods and strains

*C. elegans* were maintained at 20°C on NGM plates seeded with *Escherichia coli* B 0P50, unless otherwise stated. For *Bacillus subtilis* trials the PY79 strain was used. Some RNAi experiments were performed at 25°C. Males were cultured at low density (~5 per plate) during aging trials to reduce detrimental effects of male-male interactions. For further details, see Supplemental Materials and Methods.

### Pathology measurements

Worms were mounted onto 2% agar pads and anesthetized with 0.2% levamisole. DIC images were acquired with an 0rca-R2 digital camera (Hamamatsu) and either a Leica DMRXA2 microscope or a Zeiss Axioskop 2 plus microscope, driven by Volocity 6.3 software (Improvision, Perkin-Elmer). Images of pathology were analyzed semi-quantitatively (Garigan et al. 2002; Riesen et al. 2014)(Supplemental Fig. 1A-E). For pharynx, gonad and tumor pathologies, images were randomized, examined, assigned scores of 1-5 by two independent scorers, and mean values calculated and rounded. Here 1 = youthful, healthy appearance; 2 = subtle signs of deterioration; 3 = clearly discernible, mild pathology; 4 = well developed pathology; and 5 = tissue so deteriorated as to be barely recognizable (e.g. gonad completely disintegrated), or reaching a maximal level (e.g. large tumor filling the entire body width). Intestinal atrophy was quantified by measuring the intestinal width at a point posterior to the uterine tumors, subtracting the lumenal width and dividing by the body width. Yolk accumulation was measured by dividing the area of yolk pools with the area of the body visible in the field of view through a 63x lens.

### Single worm, longitudinal pathology analysis

Worms were cultured individually at 20°C. On days 4, 7, 11, 14 and 18 of adulthood, each worm was imaged individually by DIC microscopy (Supplemental Fig. 1F,G). For imaging, microscope slides were prepared by taping two coverslips on the slide, at each edge, leaving an empty space in the middle for the agarose pad. The worm was then placed on a 2% agarose pad on the slide. Another coverslip was then placed on top, but resting on the two side coverslips, to reduce the pressure of the coverslip onto the worm. The slide was then placed on a PE120 Peltier cooling stage (Linkam Scientific) set to 4°C. Within minutes of cooling the nematodes ceased to move, and images were taken at 630x magnification using a Zeiss Axioskop microscope. After imaging, each worm was carefully recovered by pipetting 20μL of M9 buffer between the top coverslip and the agar pad. The coverslip was then gently removed and the worm picked onto an NGM plate. Images of pharynxes, distal gonads, uterine tumors, yolky pools and intestinal atrophy were analyzed as described above. The lifespan of each nematode was then measured.

### Bodipy staining

This was performed as described (Klapper et al. 2011), except that worms were manipulated in 15μ? droplets held in parafilm micro-wells. Briefly, animals were washed 2x by transferring them successively into two droplets of M9 using a platinum wire pick. They were then transferred to a drop of 2% paraformaldehyde solution for 15-20min and frozen/thawed 3x at −80°C/room temperature (RT). Animals were then carefully washed 3x by transferring them into successive M9 droplets. They were then transferred to μg/mL BODIPY 493/503 (Invitrogen) in M9 solution for 1-2hr at RT in darkness. The worms were finally washed 3x in M9 droplets and mounted for imaging (488nm Exc./505-575nm Em.) on a Zeiss LSM710 confocal microscope.

### Lipidomic analysis

Lipidomic analysis was performed on about 3000 to 10000 individually picked worms per sample, using a Thermo Orbitrap Elite mass spectrometer. For technical details see Supplemental Materials and Methods.

### Lifespan measurements

These were performed at 25°C unless otherwise stated. Worms were either transferred daily during the reproductive period, or transferred at L4 stage to plates supplemented with 15μM FUDR to block progeny production. Animals that died from internal hatching were censored. For *atg-13* RNAi lifespan trials, blind scoring was performed.

### Quantification and statistical analysis

For pathology measurements, the Student’s t test was used. ANOVA and two-way ANOVA with Bonferroni correction were applied to multiple comparisons. Correlations from single worm, longitudinal pathology analysis were analyzed using the Spearman Rank test and linear regression analysis. Benjamini-Hochberg corrections were used for multiple comparisons. For survival assays, statistical significance was estimated using log rank and Wilcoxon statistical tests executed using JMP 11 software. Unless stated otherwise, three independent trials with N>10 in each trial were used. No statistical methods were used to predetermine sample size. The experiments were not randomized. All graphs display mean values and all error bars depict s.e.m.

### Data and software availability

The raw data that support the findings of this study are available from the corresponding author upon request. Analyzed lifespan and lipidomics data are available in supplementary tables 1 and 2.

## Acknowledgments

We thank the Caenorhabditis Genetics Center for providing many of the strains used in this study. Additional worm strains were kindly provided by A. Meléndez (C.U.N.Y.), L. Temmerman (K.U. Leuven) and H. Zhang (S.K.L.B., Beijing). This work was supported by the European Commission (FP6-518230, IDEAL), a Wellcome Trust Strategic Award (098565/Z/12/Z), and a BLS start-up fund award from Lancaster University to AB. We would also like to thank I. Bjedov, E. Nishida, L. Partridge and members of the Gems laboratory for helpful discussion, and F. Cabreiro, M. Hansen, J. Regan and J. Tullet for comments on the manuscript.

## Author contributions

A.B., M.E. and D.G. conceived and designed the study. C.A., A.B., M.E., A.F.G., D.G., T.S., S.S. and Q.Z. designed and/or performed the experiments with contributions from S.B.-N., Y.G., A.L.P., E.T. and M.J.W.. A.B., M.E. and D.G. wrote the manuscript.

## Supplemental discussion

### By promoting gut-to-yolk biomass conversion, autophagy promotes intestinal pathology

Our findings suggest that autophagy promotes intestinal atrophy and yolk steatosis. This was not expected as the role of autophagy in aging is usually viewed as a protective one, consistent with the molecular damage theory: by removing damaged cellular components, autophagy helps maintain the cell in a youthful state (Gelino and Hansen 2012). Consistent with this, inhibition of autophagy can reduce extended longevity in *C. elegans*, e.g. when induced by mutation of *daf-2* (Meléndez et al. 2003), dietary restriction (Hansen et al. 2008; Gelino et al. 2016), germline loss (Lapierre et al. 2011), brief heat-stress (hormesis) and *hsf-1* over-expression (Kumsta et al. 2017). Moreover, it has been proposed that autophagy is increased in *daf-2* mutants, which in part drives their increased longevity (Meléndez et al. 2003; Gelino and Hansen 2012; Chang et al. 2017). Yet our findings suggest that reduction of autophagy in *daf-2* mutants protects against senescent pathogenesis. How could such different claims be reconciled?

The idea that autophagy might promote rather than suppress senescent changes is not in itself contentious. Autophagy can enhance as well as inhibit the development of pathologies, including senescent ones (Shintani and Klionsky 2004; Kang and Avery 2008). For example, production of the senescence-associated secretory phenotype (SASP) proteins in mammalian cells is promoted by autophagy (Narita et al. 2011), there playing a role somewhat similar to that in *C. elegans* yolk production proposed here.

Inhibition of autophagy in *C. elegans* has been observed either to reduce or increase lifespan, or to have no effect (Meléndez et al. 2003; Hashimoto et al. 2009; Gelino and Hansen 2012; Wilhelm et al. 2017) (this study). We postulate that the pathologies that limit life vary with culture conditions and genotype (Fig. 1A), and between species. Thus, the effect on lifespan of inhibiting autophagy will depend on whether life-limiting pathologies are inhibited or enhanced by autophagy. For example, we found that *atg-13* RNAi consistently increased lifespan at 25°C but not at 20°C, despite ameliorating senescent pathologies at both temperatures (Fig. 3I, Supplemental Fig. 3H, Table 2). In another study performing RNAi on 14 autophagy genes and where increased lifespan was consistently seen (Hashimoto et al. 2009), a high concentration of 5-fluoro-2-deoxyuridine (FUDR) was used. Given that intestinal autophagy protects against bacterial infection (Jia et al. 2009), a possibility is that FUDR can alleviate life-limiting infection by co-cultured *E. coli* that autophagy protects against; autophagy also appears to protect against end of life loss of intestinal barrier function (Dambroise et al. 2016; Gelino et al. 2016).

Autophagy may also exert life-shortening and life-extending effects at different times in *C. elegans* life history. Reduced autophagy during development can impair health (Hashimoto et al. 2009; Zhang et al. 2015), while late-life knock-down of autophagy genes (including *atg-13*, studied here) can increase lifespan (Wilhelm et al. 2017). Effects of autophagy knockdown on lifespan may also depend upon severity of knockdown. The adult hermaphrodite intestine exhibits relatively high levels of autophagy (Chapin et al. 2015) and protein turnover (Dhondt et al. 2017); we speculate that high intestinal autophagy levels that maximize yolk production exceed homeostatic requirements, such that partial inhibition of autophagy can improve late-life health, while full abrogation impairs essential housekeeping functions and reduces health. More generally, where wild-type levels of autophagy promote life-limiting pathology, moderate inhibition in autophagy may extend lifespan in the wild type, but shorten it in a long-lived mutant in which autophagy levels are already optimally reduced.

### Does IIS increase or decrease intestinal autophagy?

The status of autophagy in *daf-2* mutants remains unclear. Reliable means to directly measure autophagic activity in *C. elegans* have yet to be developed (Zhang et al. 2015; Chang et al. 2017). For example, autophagosome numbers in *daf-2* mutants are increased in the larval hypodermis (Meléndez et al. 2003) and in the adult intestine (Chang et al. 2017). This could reflect either increased autophagosome production (increased autophagy) or a slowing-down in autophagosome consumption (decreased autophagy) (Zhang et al. 2015; Chang et al. 2017). However, autophagy does not appear to be blocked in the *daf-2* mutant intestine (at least, not entirely) suggesting increased autophagy (Chang et al. 2017). Another broad indicator of autophagic activity is protein turnover rate, although this is also a function of ubiquitin/proteasomal activity. For example, aging *C. elegans* show a decline in both autophagy (Chang et al. 2017) and protein turnover (Depuydt et al. 2016; Dhondt et al. 2017). However, *daf-2* mutants show major reductions in both protein synthesis (Depuydt et al. 2013; Stout et al. 2013) and turnover (Depuydt et al. 2016; Dhondt et al. 2016; Visscher et al. 2016). Thus, there is evidence of both increased and decreased autophagy in *daf-2* mutants. Our findings support the latter view, and suggest a mechanism by which autophagy acts to promote senescent pathogenesis (gut-to-yolk biomass conversion). Given that IIS strongly promotes yolk production (Depina et al. 2011), intestinal atrophy could result from the combined effects of IIS-promoted dominance of the translational machinery by vitellogenin synthesis, constitutively high levels of intestinal autophagy (Chapin et al. 2015; Dhondt et al. 2017), and yolk export (Kimble 1983). Another possibility is that IIS promotes TOR-activated, coupled protein synthesis and autophagy (Narita et al. 2011). In conclusion, this discussion provides a new perspective on the role of autophagy in *C. elegans* aging, which we hope will usefully inform future investigations of the topic.

## Supplemental Materials and Methods

### Culture methods and strains

*C. elegans* were maintained in standard conditions (Brenner 1974), at 20°C on NGM plates seeded with *Escherichia coli* OP50, unless otherwise stated. For *Bacillus subtilis* trials the PY79 strain was used. Some RNAi experiments were performed at 25°C. Males were cultured at low density (~5 per plate) during aging trials to reduce detrimental effects of male-male interactions (Gems and Riddle 2000b). The following strains were used: N2 (wild type N2 male stock, N2 CGCM) (Gems and Riddle 2000a); CB3168 *him-1(e879); mab-3(e1240)* (Shen and Hodgkin 1988), CF80 *mab-3(mu15); him-5(e1490),* DH26 *rrf-3(b26),* DR466 *him-5(e1490)*, DR1563 *daf-2(e1370)*, DR1572 *daf-2(e1368)*, GA1500 *bIs1[pvit-2::vit-2::GFP + rol-6(su1006)]*, HZ1683 *him-5(e1490); atg-2(bp576),* HZ1685 *atg-4.1(bp501)*, HZ1688 *atg-13(bp414)*, LIU1 *ldrIs1 [pdhs-3::dhs-3::GFP + unc-76(+)]*, LSC897 *ceh-60(lst466)*, LSC903 *ceh-60(lst491)*, LSC904 *lrp-2(lst464),* RT311 *unc-119(ed3); pwIs69 [vha6p::GFP::rab-11 + unc-119(+)],* RT476 *unc-119(ed3); pwIs170 [vha6p::GFP::rab-7 + Cbr-unc-119(+)]*, RT525 *unc-119(ed3); pwIs206 [vha6p::GFP::rab-10 + Cbr-unc-119(+)]*, RT1315 *unc-119(ed3); pwIs503 [pvha-6::mans::GFP + Cbr-unc-119(+)]*, VC893 *atg-18(gk378)*, VP303 *rde-1(ne219); kbEx200 [pnhx-2::rde-1]*. The following strains were created for this study: GA1711 *wuEx272 [pvit-6::vit-6::mCherry + rol-6]*, GA1715 *wuEx280[rol-6]*, GA1729 *atg-13(bp414); wuEx292[pges-1::atg-13 + rol-6]*, GA1730 *atg-13(bp414); wuEx293[pges-1::atg-13 + rol-6],* GA2100 *bIs1[pvit-2::vit-2::GFP + rol-6]; wuEx277[pvit-6::vit-6::mCherry + rol-6]*.

### Generation of transgenic strains

Constructs were made using PCR fusion using primers as follows. For the *vit-6::mCherry* construct, the 6,740 bp *vit-6* promoter and genomic region, excluding the stop codon, was amplified using F: 5’-TTCTTCTTTCGGTGGCTCTG-3’ and R: 5’-CTTCTT CACCCTTT GAGACCAT AT AGT CGAACTT GT CGCACT −3’. mCherry was amplified using F: 5’-ATGGTCTCAAAGGGTGAAGAAG-3’ and R: 5’-GATGGCGATCTGATGACAGC-3’.

The 3’UTR was amplified using F: 5’-CTACCTCTTCTTCACAATCATACAC-3’ and R: 5’-ACTGTAGAAGTGAACTCTGTG-3’. The *vit-6* fragment was fused to mCherry using F: 5’-TGGAGACACAATAGAAGTCG-3’ and R: 5’-GTGTATGATTGTGAAGAAGAGGTAGCTACTTATACAATTCATCCATGCCAC-3’. The fused fragment was further fused with the 3’UTR using F: 5’-ATTCCACAGAAAGGATTGCAC-3’ and R: 5’-ATGCCGAGTTGTTTGAATTG-3’. For the *atg-13* intestinal rescue, the *ges-1* promoter was amplified using F: 5’-TTGTCTATTGGTATGGCTGC-3’ and R: 5’-GTACGTGTCGTACTCATTTACCATACAAGGAATATCCGCATCTG-3’. The *atg-13* 2,300 bp genomic region was amplified using F: 5’-ATGGTAAATGAGTACGACACGTAC-3’ and R: 5’-TGCAAGACTTCTGAGCAATG-3’. The two fragments were fused using primers F: 5’-GCGCTACCAATAAGGCTAAG-3’ and R: 5’-GAGCAATGTCGCAATGGAAAG-3’. *vit-6::mCherry* was microinjected at 1μg/μL with 100μg/μL *rol-6* coinjection marker to generate GA1711. GA1711 was crossed with GA1500 to obtain GA2100. *pges-1::atg-13* was injected into *atg-13(bp414)* at 80μg/μL with 20μg/μL *rol-6* coinjection marker. Two independent lines, GA1729 and GA1730, were used for experiments.

### Electrophoretic analysis of yolk proteins

Yolk protein levels were quantified by running worm protein extracts on PAGE gels and then staining with Coomassie blue dye as described (Depina et al. 2011). 20 worms were picked into 25μL of M9 buffer and frozen at −80°C. Samples where then thawed, and 25μL of 2x Laemmli sample buffer (Sigma) added. Samples were incubated at 70°C, vortexed continuously for 15 min, incubated at 95°C for 5 min and at 6,000 rpm for 15 min. Samples were loaded onto Criterion XT precast gels 4-12% Bis-Tris (Bio-Rad), using XT MOPS (Bio-Rad) as a running buffer, and stained and destained following standard protocols. Gels were analyzed using ImageQuant LAS-4000 (GE Healthcare). Protein band identification was based on published data (Kimble 1983; Depina et al. 2011). Yolk proteins were normalized to myosin and the ratio of actin to myosin was estimated to assess the reliability of myosin as a standard for normalization (Depina et al. 2011).

### Yolk protein proportion measurements

Two approaches were taken to estimate the proportion of total worm protein that is yolk protein: using densitometry or reference to protein standards of known concentration. For the densitometry-based approach, the density of individual vitellogenin bands (YP170, YP115 and YP88), and of the entire lane, of protein gels stained with Coomassie Blue were measured. The density of a whole lane (all protein) was set as 100%, and the percentages of the vitellogenin bands were calculated proportionally. The optimal threshold for densitometric reading was that which was just sufficient to exclude background from regions between gel lanes. For the protein standard-based approach, known amounts of protein standard were run alongside worm protein samples and gels stained with Coomassie Blue. The density of bands obtained from standard protein was used to construct a standard curve. The intensity of vitellogenin bands (YP170, YP115 and YP88) from the samples was then compared to the standard curve, to calculate the amounts of vitellogenin protein present. To estimate the proportion of yolk proteins to total protein, the estimated amount of yolk protein was compared with total protein content data obtained from protein quantification using BCA.

### Total nematode protein measurement

This was performed using bicinchoninic acid (BCA). Samples were prepared by adding 250μL of Cell Lytic M buffer (Sigma) containing 1:1000 protease inhibitor cocktails. Samples were then mixed and sonicated using a Bioruptor (Cosmo Bio Co., Ltd, Tokyo, Japan) for 8 min at 30 sec intervals. Samples were centrifuged at 4°C at 6000 rpm for 15 min. The BCA method was performed in a 96-well plate, with each well containing 200μL of testing solution and 25μL sample or bovine serum albumin (BSA) standards. The plate was mixed gently and incubated at room temperature for 2 min and then incubated at 37°C for 30 min. Absorbance was then measured at 620 nm.

### Confocal microscopy

Bodipy stained animals were mounted on an 2.5% agarose pad between slide and coverslip in M9. 25-35μm thick Z-stacks were acquired every 0.75μm through Zeiss Plan-Apochromat 40X or 63X 1.4 immersion lenses using Zeiss LSM710 (UCL) and LSM880 (LU) confocal microscopes equipped with 405 nm, 488 nm, 568 nm and 633 nm lasers, and controlled by the Zen software package. Bodipy, VIT-2::GFP, DHS-3::GFP and MANS-2::GFP were imaged using the 488 nm laser, while VIT-6::mCherry was imaged using the 568 nm laser. Confocal image analysis of Bodipy staining focused on the int1, 2 and 3 gut cells, averaging fluorescence from 7 consecutive Z-planes that cut through the intestinal lumen. Within the ROI, segmentation in intestinal and non-intestinal areas was performed plane by plane comparing brightfield and fluorescence images across the Z-stack to delineate the intestinal limits. For DHS-3::GFP and MANS-2::GFP, scoring systems were established using a scale of 0 to 10 for DHS-3::GFP and 0 to 5 for MANS-2::GFP, with 0 representing no significant staining and 10 or 5 representing even and bright labeling. Intermediary scores correspond to various degrees of signal amount (area), density, heterogeneity and sharpness. The results shown are the average of two rounds of scoring for each dataset. Each confocal microscopy experiment combines 2 to 4 independent replicates of 5 to 30 worms for each genotype/time point combination. As fluorescence levels and dynamics vary between worms and slides, fluorescence intensity cannot be reliably related to lipid amounts, which is why lipid organelle area was chosen over fluorescence intensity as quantitative measure. All fluorescence quantifications were performed on raw images while illustrations in supplemental figures were saturated to enable easy visualization.

### Electron microscopy

30-40 adult worms per condition were pre-fixed in a drop of 2% low-melting point agarose solution (kept below 30°C on a heat block) containing 2.5% glutaraldehyde, 1% paraformaldehyde in 0.1M sucrose, 0.05M cacodylate, and 0.02% levamisole. Worms were moved to a dissecting scope at room temperature and quickly aligned as the agarose set, and then cut in half using a razor blade. Half worms were realigned upon addition of another drop of melted agarose, and the gelled block was then trimmed before transferring into the fixative solution (2.5% glutaraldehyde, 1% paraformaldehyde in 0.1M sucrose, and 0.05M cacodylate) on ice. The agarose block was then stained and processed as described (Shaham 2006) (protocol 8), adjusting timings to account for slower diffusion through the agarose block. Serial 1μ? sections were taken for light microscopy inspection, and ultra-thin (70-80 nm) sections were cut at the region of interest using a diamond knife on a Reichert ult.ramicrot.ome. Sections were collected on slot grids and stained with lead citrate before viewing using a Joel 1010 transition electron microscope. Images were captured with a Gatan Orius camera and Gatan imaging software, and then exported in TIFF format.

### Total triacylglyceride quantification

Biovision’s Triglyceride Quantification Kit (Mountain View, CA) was used to assay for triacylglyceride content. Animals were aged and counted before harvest. Samples contained approximately 300 animals in 100μL Biovision assay buffer and were frozen in liquid nitrogen and thawed 100°C 3 times, followed by sonication to break the cuticle, followed for sonication. Lipids were extracted in glass tubes with the Folch method and reconstituted in 100μL assay buffer. Triacylglycerides were measured using the Biovision's Triglyceride Quantification Kit protocol.

### Lipid extraction and analysis

Worms were counted and collected in M9 buffer. This was followed by washing: samples were centrifuged at 1000 rpm for 3min, allowing the worms to collect at the bottom of the tube but the bacteria to stay in suspension. As much as possible of the supernatant was removed without disturbing the worm pellet and fresh M9 was then added. The process was repeated a few times, until the supernatant was clear and free of bacteria. After a final wash in PBS, the worm pellet was resuspended in 1mL PBS and frozen at −80°C. The samples were thawed while vortexing at room temperature and transferred to 10mL silanized glass tubes using silanized glass pipettes. 1mL methanol and 2mL chloroform were added to each sample and mixed with a vortex. 40μL of lipid standard mixture (12:0/12:0/12:0-triacylglycerol (TG; 800 ng)) were used to spike each sample. After thorough mixing, samples were subjected to modified Folch extraction. The lipid extracts were dissolved in 150yL chloroform in silanised glass sample vials. lyL of sample were injected for positive ion lipids analysis and another lyL was injected for negative ion lipids analysis. In brief, different classes of lipids were separated on normal phase Type C silica gel column (150×2.1mm, 4ym, 100A, MicoSolv Technology) with hexane/dichloromethane/chloroform/methanol/acetanitrile/water/ethylamine solvent gradient elution based on the polarity of head group on Shimadzu Prominence HPLC system. Individual lipid species were identified and semi-quantified with high resolution (240 k at m/z 400)/accurate mass analysis (mass accuracy <5ppm) on Thermo Orbitrap Elite mass spectrometer.

### Intestinal autophagy assays

These were performed using strain DLM3 *ttTi5605 II; unc-119(ed3) III; uwaEx2 [vha-6p::CERULEAN-VENUS::lgg-1 + unc-119(+)]*, as described (Chapin et al. 2015). Animals were maintained on *E. coli* HT115 and the test group subjected to *atg-13* RNAi from L4 onwards (20°C). mFP/dFP ratio provides a relative measure of autophagy (specifically, autolysosome formation).

### Mating and male scent exposure tests

To test effects of mating on pathology, single hermaphrodites were mated with 3 males. On day 3, males were removed and mated hermaphrodites subsequently identified by the presence of male progeny, and then maintained for pathology analysis. To test effects of male scent, 60mm NGM plates were conditioned with males for 2 days. Males were then removed and hermaphrodites added. Plates were either conditioned with 150 males and then 30 hermaphrodites added, or with 60 males and then 30 hermaphrodites added. In a further protocol, 35mm NGM plates were conditioned with 30 males to which 30 hermaphrodites were added.

---

**Supplemental Figure 1.**
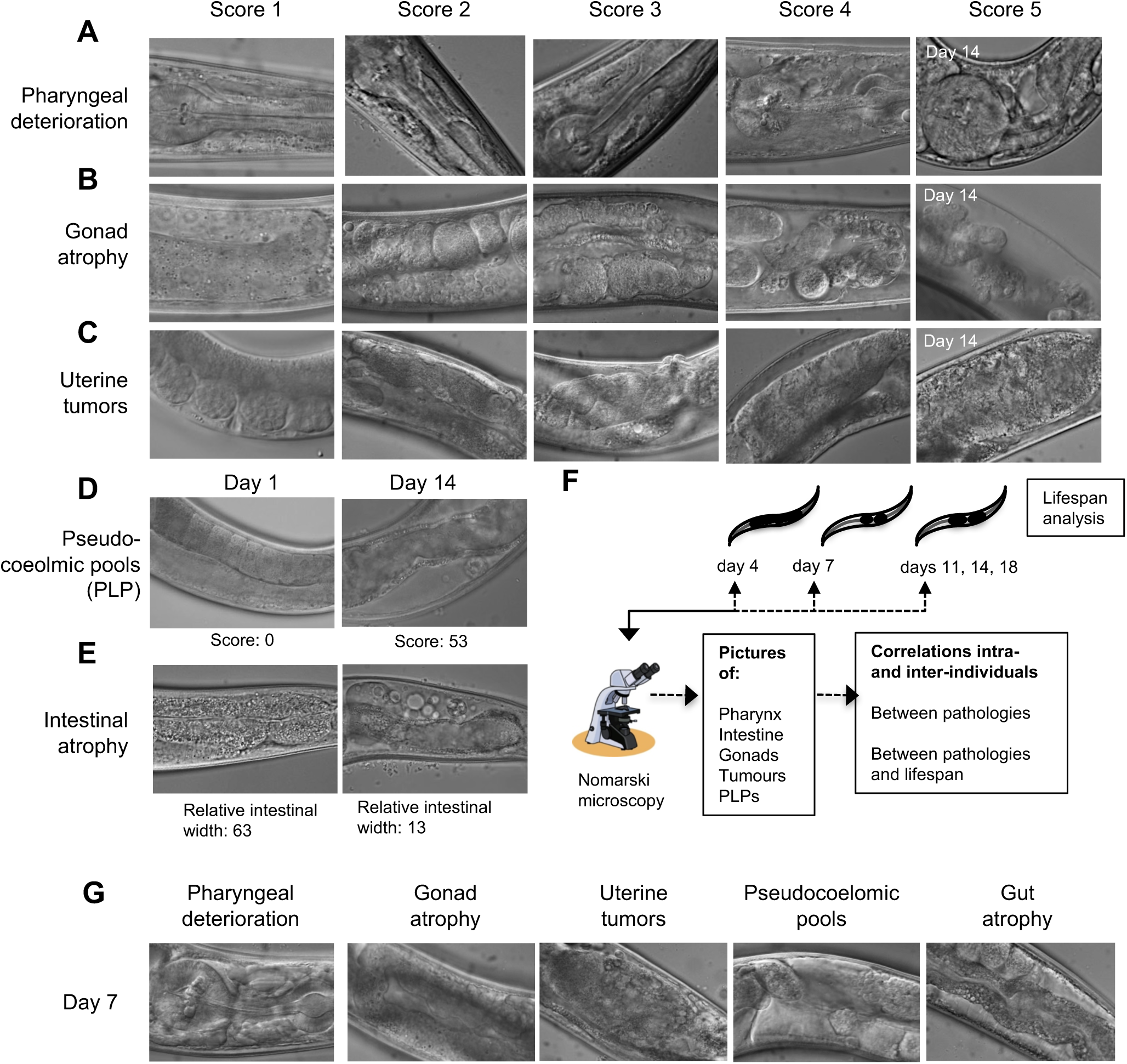
Semi-quantitative estimation of severity of senescent pathology. (*A-C*) For pharynx (*A*), gonad (*B*) and uterine tumor (*C*) pathologies, images were randomized, and given scores of 1-5. Here 1 = a youthful, healthy appearance; 2 = showing subtle signs of deterioration; 3 = clearly discernible, low level pathology; 4 = well developed pathology; and 5 = tissue so deteriorated as to be barely recognizable (e.g. gonad completely disintegrated), or reaching a maximal level (e.g. large tumor filling the entire body diameter). (*D, E*) Pseudocoelomic pool formation (yolk accumulation) (*D*) and intestinal atrophy (*E*) were measured by dividing the total area of yolk pools with the area of the body visible in the field of view. (*F*) Experimental approach for measuring correlation between severity of senescent pathologies and lifespan, by combining serial imaging of pathology in individual nematodes, and then measuring their lifespan. (*G*) Examples of images of pathology at one of the time points examined (day 7).

**Supplemental Figure 2.**
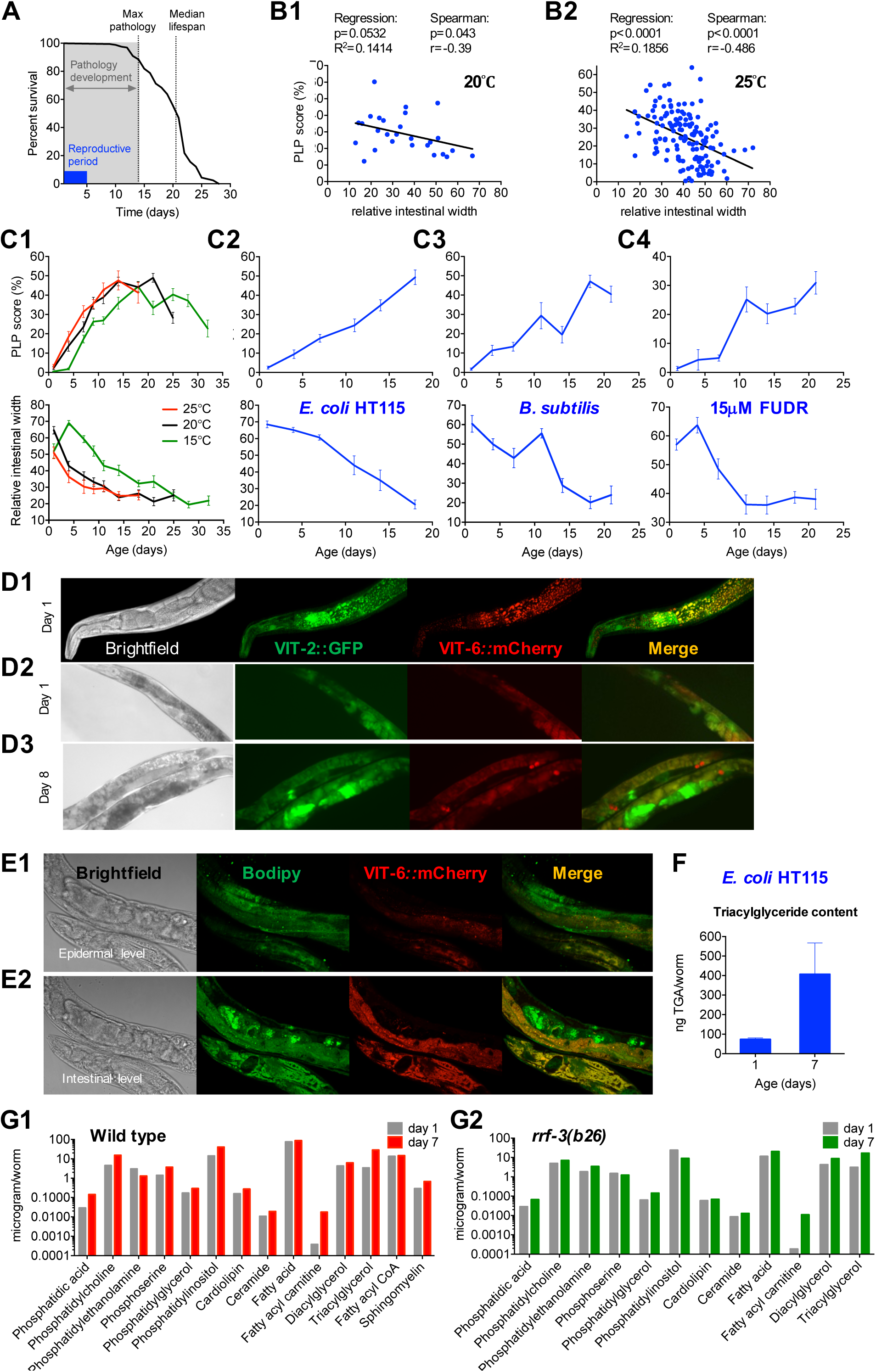
Senescent PLP accumulation and intestinal atrophy are robustly correlated and coincide with an age-shift in lipid composition. (*A*) Wild type adulthood at 20°C is characterized by a 4-5 day self-reproduction period, 14-days of senescent pathology development, and a 20 day mean lifespan. (B) CP accumulation score and intestinal atrophy correlate at the individual worm level at 20°C without FUDR (*B1*) and at 25°C with FUDR (*B2*). (*C*) Co-development of CP accumulation (upper panels) and intestinal atrophy (lower panels) with age occurs at all temperature tested (*c1*), on *E. coli* K12 diets (*C2*), on a *Bacillus subtilis* diet (*C3*), and in presence of FUDR (*C4*). (*D*) Columns from left to right show Nomarski, VIT-2::GFP, VIT-6::mCherry and merge of VIT-2::GFP and VIT-6::mCherry, respectively. Day 1 adults show VIT-2::GFP and VIT-6::mCherry in the intestine (200X magnification) (*D1*). Day 1 adults show VIT-2::GFP and VIT-6::mCherry in the intestine and embryos (100X) (*D2*). Day 8 adults show VIT-2::GFP and VIT-6::mCherry in the intestinal droplets and CPs (100X) (*D3*). VIT-2::GFP was also observed in the tumors and VIT-6::mCherry in the coelomocytes. (*E*) Columns from left to right show Nomarski, Bodipy, VIT-6::mCherry and merge of Bodipy and VIT-6::mCherry, respectively. Day 10 adults, epidermal level, show Bodipy staining of lipid droplets within muscles (200X) (*E1*). Day 10 adults, intestinal level, show yolky lipid within and outside of the intestine (200X) (*E2*). (*F*) Quantification of triglyceride (TAG) levels. TAG content increases 5-fold between day 1 and day 7 in Wildtype hermaphrodites maintained on *E. coli* HT115. (*G*) Mass spectroscopy analysis of age changes in lipid profiles. In wild type at 20° C TAG content increases 8.2 fold (*G1*). In sterile *rrf-3(b26)* mutants at 25° C TAG content increases 5.0 fold (*G2*).

**Supplemental Figure 3.**
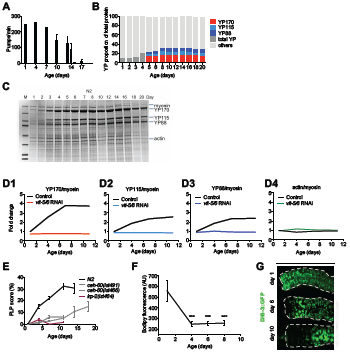
Labeling of yolk proteins and lipids in aging hermaphrodites. (*A*) Quantification of pharyngeal pumping. Feeding rate declines with age. (*B*) Quantification ofYP proportion of total protein using protein standards. YP protein proportion increases from less than 10% on day 1 to ~35% on day 8. (*C*) Representative Coomassie stained gel showing age-increase in yolk protein (YP) levels in N2 hermaphrodites; cf. Fig. 1h. (*D*) *vit-5,-6* RNAi abrogates age increase in yolk protein (YP) accumulation (*D1-D3*). (*E*) Reduced accumulation of CPs in *ceh-60* and *lrp-2* mutants. (*F*) Quantification of intestinal lipid content using Bodipy. Bodipy staining in the intestine decreases with age. Trials conducted at 25 °C with 15μ? FUDR. (*G*) Images of intestinal lipid droplets using DHS-3::GFP. Numbers of lipid droplets showing DHS-3::GFP fluorescence decreases with age.

**Supplemental Figure 4.**
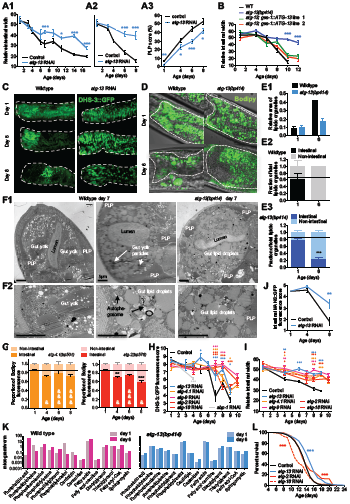
Impact of autophagy inhibition on visceral pathologies in aging hermaphrodites. (*A*) Adult-limited *atg-13* RNAi suppresses intestinal atrophy and PLP accumulation; 25 °C. Relative intestinal width is suppressed at 20 °C, no FUDR (*A1*), and at 25°C, 15μ? FUDR (*A2*). Suppression of PLP accumulation at 25°C, 15μ? FUDR (*A3*). (*B*) Intestine-specific rescue of *atg-13(bp414)* restores intestinal atrophy to wildtype levels; 25°C, no FUDR. (*C*) Labeling of intestinal lipid droplets using DHS-3::GFP in control and *atg-13* RNAi animals. (D) Bodipy staining of neutral lipids in control and *atg-13* RNAi animals. (*E*) Quantification of lipidic organelles using TEM. Age-related increase in lipidic area is suppressed by *atg-13(bp414)* (*E1*). Relative intestinal lipidic organelle content is preserved during aging in *atg-13(bp414)* compared to wildtype (*E2, E3*). (*F*) Transmission electron microscopy (TEM) images at 3,000X (*F1*) and 10,000X (*F2*) of wildtype and *atg-13(bp414)* worm sections on day 7 of adulthood. *atg-13(bp414)* worms have reduced PLPs and increased intestinal lipid droplets compared to wildtype. (*G*) Bodipy staining indicates that abrogation of *atg-4.1(bp501)* or *atg-2(bp576)* retards senescent redistribution of lipid (* comparison to day 1, & comparison to age-matched control). (*H, I*) Adult-limited inhibition of autophagy *[atg-2* RNAi, *atg-4.1* RNAi, *atg-9* RNAi, *atg-13* RNAi, *atg-18* RNAi] delays DHS-3::GFP-tagged intestinal lipid droplet reduction (H) and intestinal atrophy (**I**). (*J*) Quantification of Golgi abundance using MANS::GFP. Age-related decrease in Golgi abundance is suppressed by *atg-13* RNAi. (*K*) *atg-13(bp414)* blocks the age-associated lipidomic shift occurring between day 1 and day 6 of adulthood; 25°C, 15μm FUDR. (*L*) Adult-limited *atg-2* RNAi or *atg-13* RNAi increase while *atg-18* RNAi decreases lifespan; 25°C, 15μm FUDR. */& p<0.05, **/&& p<0.01, ***/&&& p<0.001 (* comparison to age-matched control).

**Supplemental Figure 5.**
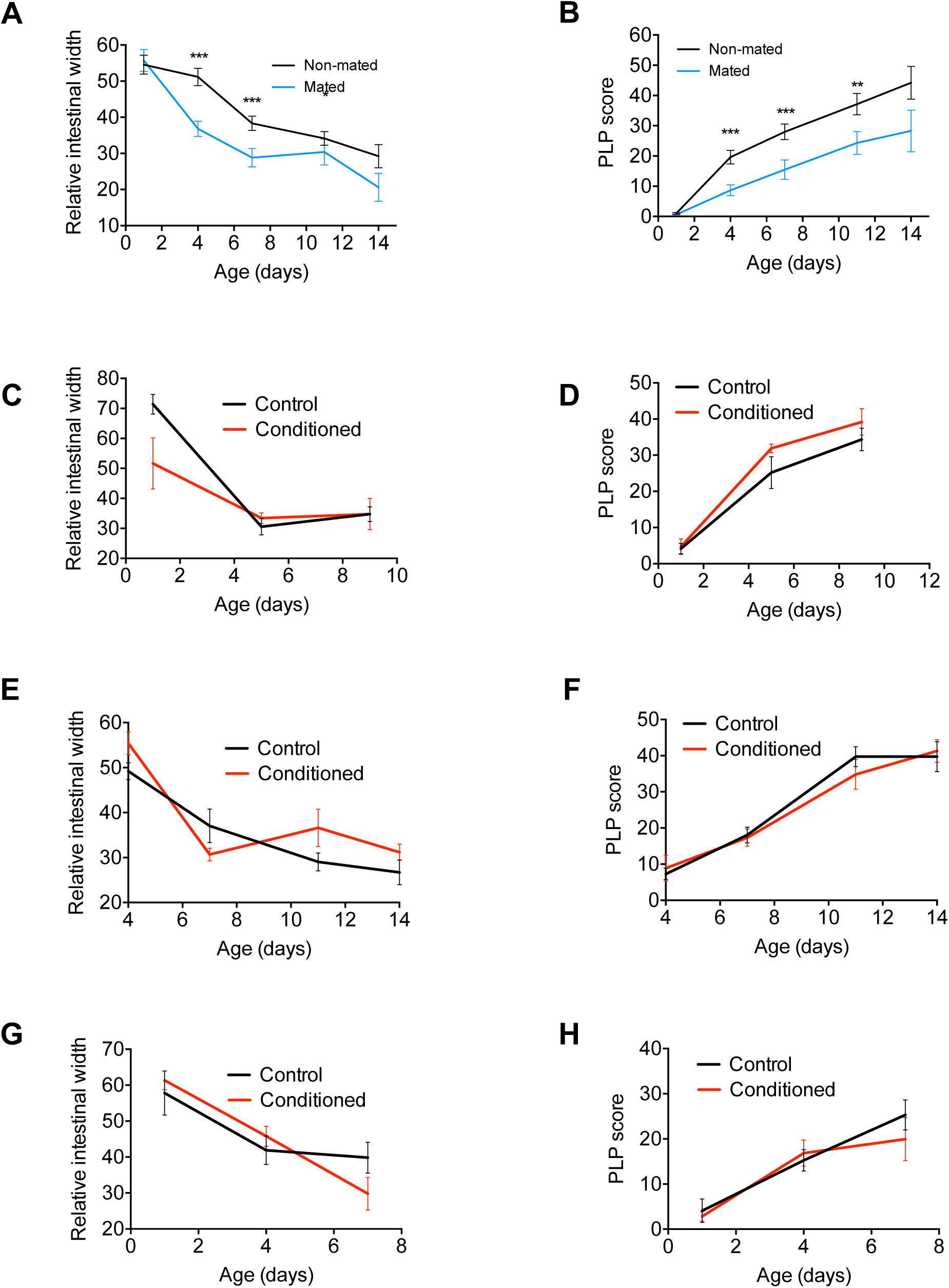
Effects of mating and exposure to males on visceral pathology. (*A, B*) Effects of mating on intestinal pathology (*A*) and yolk steatosis (*B*) Each hermaphrodite was mated with three males for three days. (*C, D*) Effects of exposure to male scent on intestinal pathology and yolk steatosis. Conditioning experiments were performed with different ratios of conditioning males and assayed hermaphrodites. 150 males:30 hermaphrodites (*C-D*). 30 males:30 hermaphrodites (*E-F*). 60 males:30 hermaphrodites on 40mm plates (*G-H*).

**Supplemental Figure 6.**
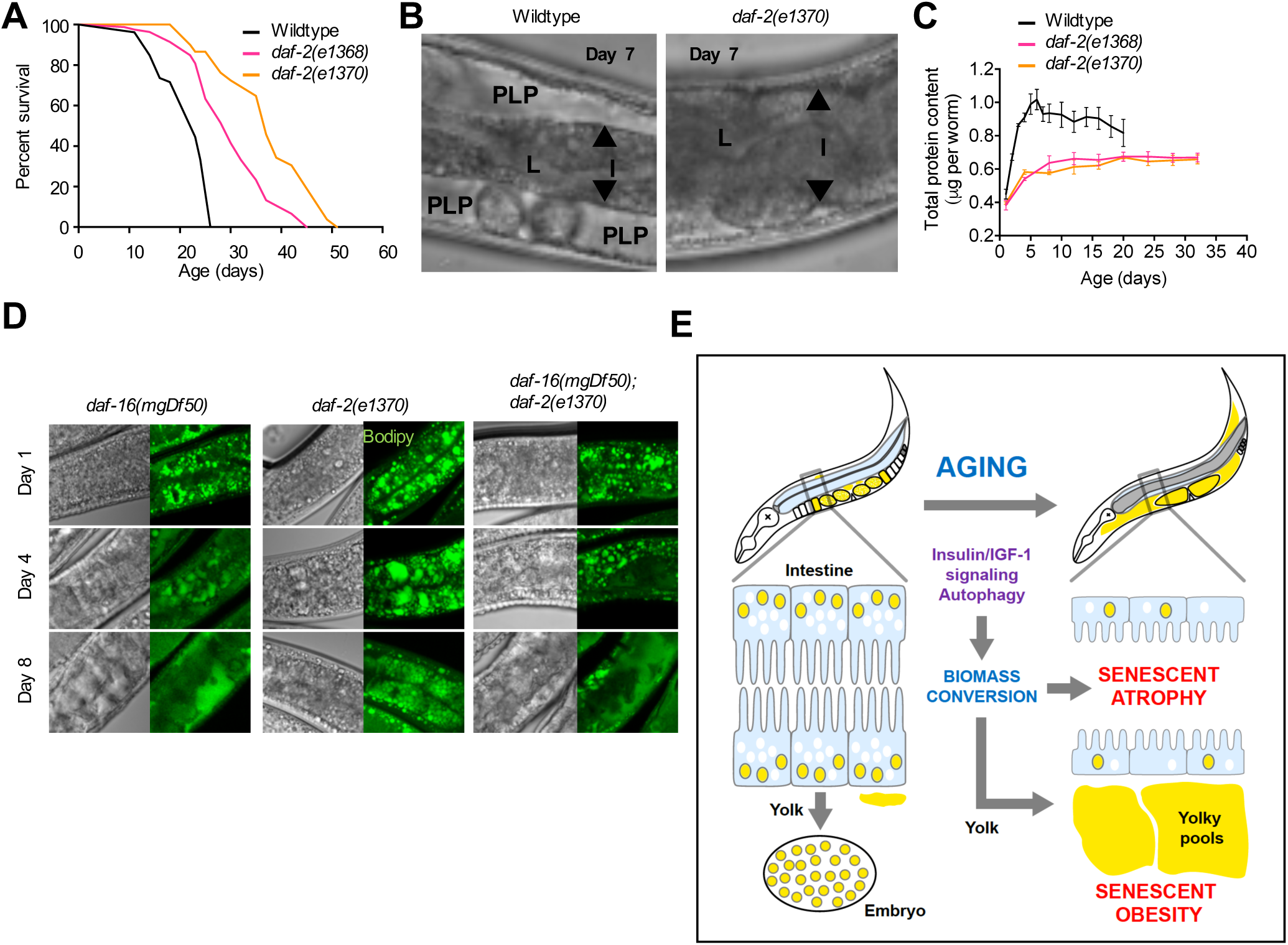
Gut-to-yolk biomass conversion results in gut atrophy and yolky pool accumulation. (*A*) daf-2(e1370) mutants (class II) live longer than *daf-2(e1368)* mutants (class I), 20°C. (*B*) Left: Representative Nomarski picture of wild-type N2 hermaphrodite mid-body at day 14 of adulthood (20°C), showing atrophied intestine and yolky pool accumulation. Right: age-matched picture of *daf-2(e1370)* mutant mid-body, showing a full-sized intestine and absence of yolky pools. (*C*) Comparison of total protein contents during aging in age-matched wild type, *daf-2(e1368)*, and *daf-2(e1370)* mutant, *daf-2* mutant protein content increases much less with age than wild type. (D) Representative pictures of gentle-fix Bodipy staining in the anterior intestine and surrounding tissues in *daf-16(mgDf50)* (left), *daf-2(e1370)* (middle), and *daf-16; daf-2* mutants (right), illustrating DAF-16-dependent preservation of intestinal fat stores and prevention of fat transfer in *daf-2* mutants at days 1, 4 and 8 of adulthood at 25°C.

**Supplemental data Table 1.**
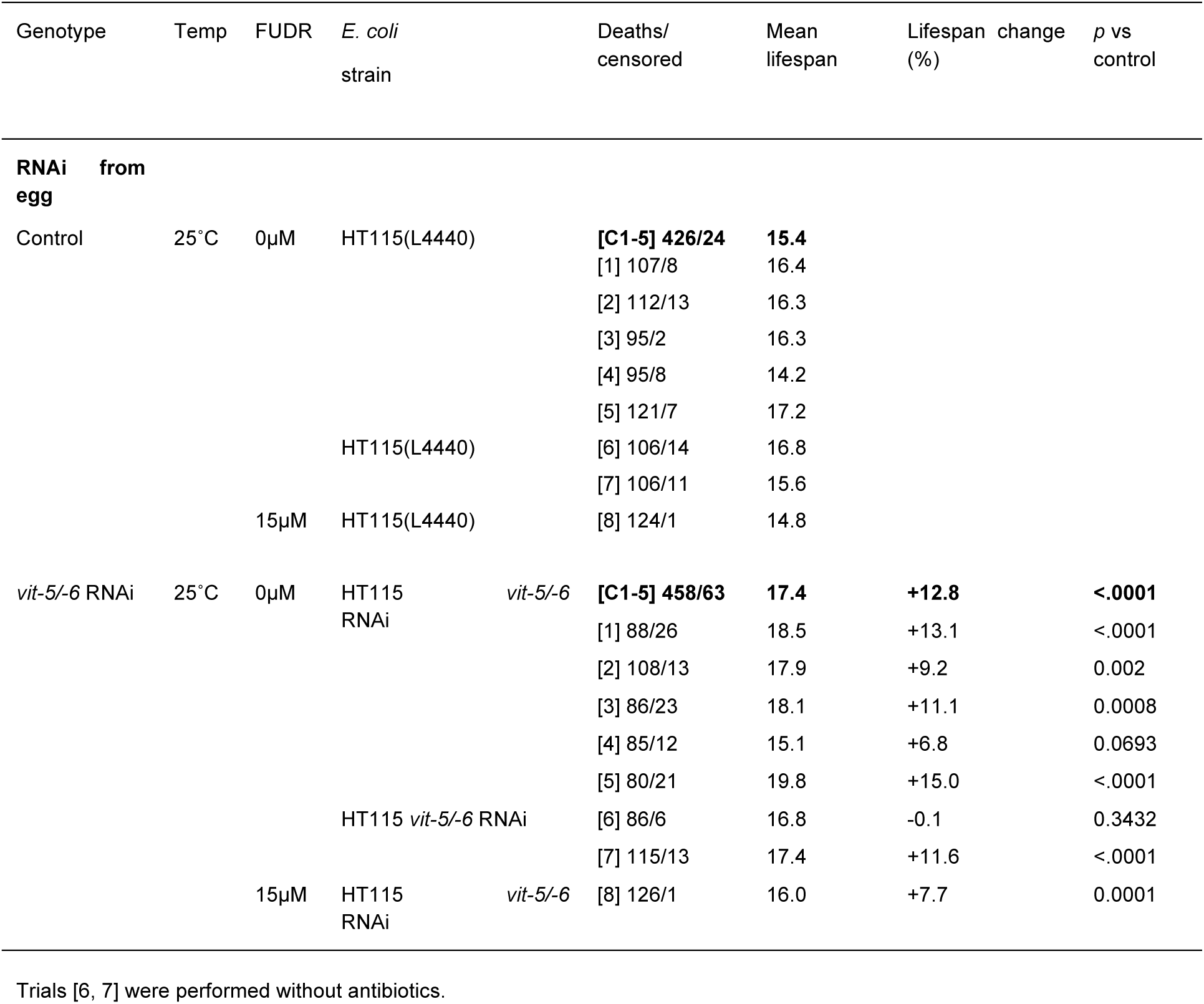
Summary statistics of lifespan trials (A) Effects of *vit-5/-6* RNAi from hatching in wildtype *C. elegans*

**Supplemental data Table 1.**
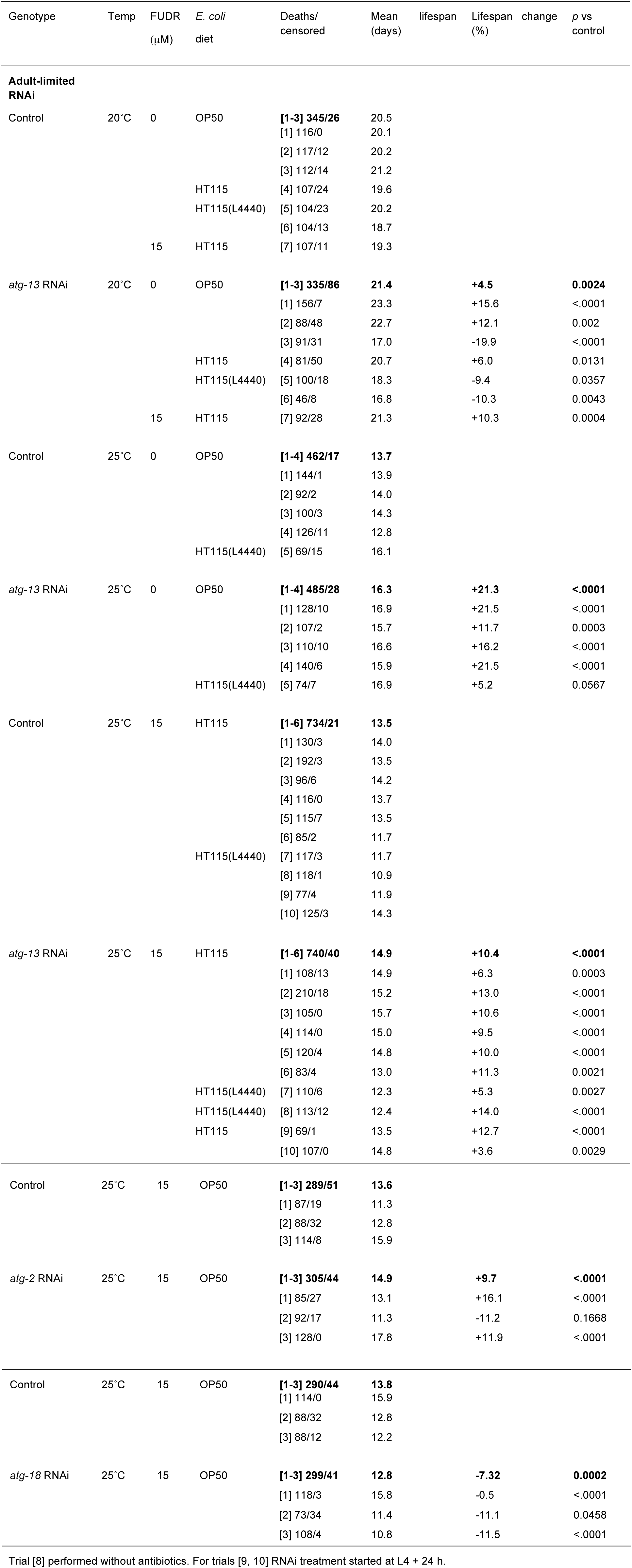
Summary statistics of lifespan trials (B) Effects of autophagy gene RNAi from adulthood in wildtype *C. Elegans*

**Supplemental data Table 1.**
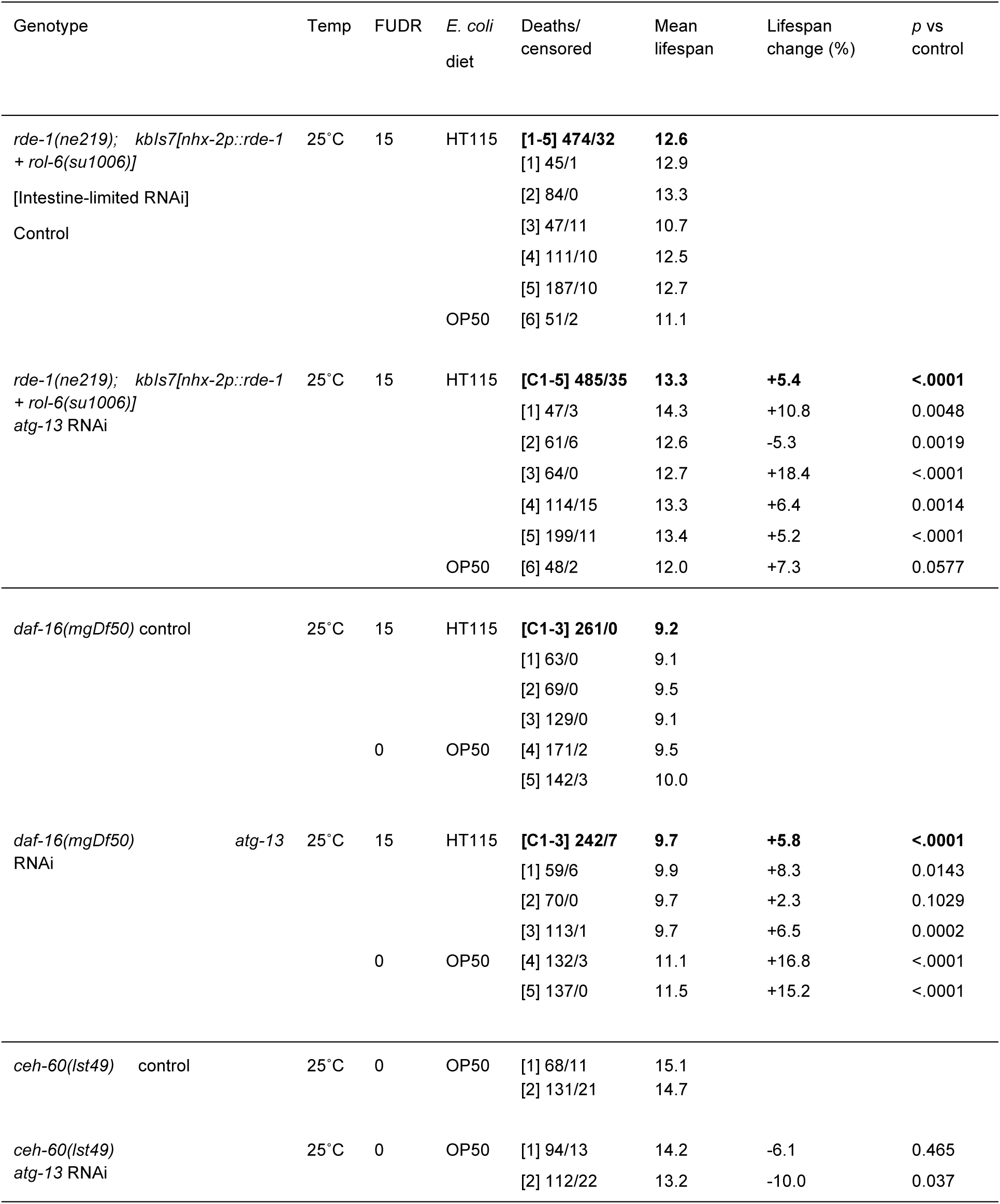
Summary statistics of lifespan trials (C) Effects of *atg-13* RNAi from adulthood in mutant *C. elegans*

**Supplemental data Table 2.** Lipid profiles of young and ageing adult wild type and *rrf-3(b26)* sterile mutant *C. elegans* hermaphrodites at 20°C, and day 1 and day 6 wild type and *atg-13(bp414)* autophagy mutant *C. elegans* hermaphrodites at 25°C with 15μM FUDR.

| Lipid species<br>(ng per worm) | 20°C no FUDR |  |  |  | 25°C 15µM FUDR |  |  |  |
| --- | --- | --- | --- | --- | --- | --- | --- | --- |
|  | Wild<br>type day<br>1 | Wild<br>type day<br>7 | *rrf-3*<br>day 1 | *rrf-3*<br>day 7 | Wild<br>type day<br>1 | Wild<br>type day<br>6 | *atg-13*<br>day 1 | *atg-13*<br>day 6 |
| Phosphatidic acid | 0.0197 | 0.1060 | 0.0287 | 0.0664 | 0.0254 | 30.4577 | 0.0280 | 0.0639 |
| Phosphatidylcholine | 1.8859 | 6.0791 | 4.9495 | 7.0140 | 1.7199 | 0.1552 | 1.6067 | 1.6504 |
| Phosphoethanolamine | 1.1001 | 1.7826 | 1.8407 | 3.4344 | 0.6872 | 2.2187 | 0.4427 | 0.6395 |
| Phosphatidylserine | 0.2397 | 0.8127 | 1.4856 | 1.2290 | 0.1925 | 0.5967 | 0.1815 | 0.2185 |
| Phosphatidylglycerol | 0.1081 | 0.1725 | 0.0629 | 0.1434 | 0.0526 | 0.3416 | 0.0349 | 0.0488 |
| Phosphatidylinositol | 9.5943 | 30.0363 | 23.8472 | 8.9514 | 4.2035 | 0.0330 | 6.8700 | 10.6244 |
| Cardiolipin | 0.1187 | 0.1681 | 0.0589 | 0.0686 | 0.0807 | 0.0287 | 0.0708 | 0.0578 |
| Ceramides | 0.0012 | 0.0029 | 0.0085 | 0.0128 | 0.0029 | 0.0036 | 0.0013 | 0.0044 |
| Fatty acids | 64.78 | 75.82 | 11.45 | 20.19 | 11.61 | 27.41 | 10.91 | 8.43 |
| Fatty acyl-carnitine | 0.0002 | 0.0131 | 0.0002 | 0.0109 | 0.0027 | 0.0040 | 0.0021 | 0.0039 |
| Diacylglycerol | 2.7751 | 4.5304 | 4.2648 | 8.7420 | 0.7787 | 1.8314 | 0.6915 | 1.1561 |
| Triacylglycerol | 1.5930 | 13.2132 | 3.0936 | 16.7488 | 1.1455 | 1.2188 | 0.9313 | 0.8023 |
| Fatty acyl-CoA | 13.479 | 14.470 | - | - | 0.290 | 1.599 | 0.183 | 0.181 |
| Sphingomyelin | 0.0637 | 0.2101 | - | - | 0.0480 | 0.1352 | 0.0536 | 0.1271 |
| Others | 4.858 | 12.967 | 12.477 | 5.577 | 1.448 | 5.007 | 2.197 | 2.254 |
| TOTAL | 100.62 | 160.39 | 63.56 | 72.19 | 22.28 | 71.04 | 24.21 | 26.26 |

- Missing values (compromised samples).

## References

Baker DJ, Childs BG, Durik M, Wijers ME, Sieben CJ, Zhong J, Saltness RA, Jeganathan KB, Verzosa GC, Pezeshki A et al. 2016. Naturally occurring p16(Ink4a)-positive cells shorten healthy lifespan. Nature 530: 184-189.

Beckman KB, Ames BN. 1998. The free radical theory of aging matures. Physiol Rev 78: 547-581.

Blagosklonny MV. 2006. Aging and immortality: quasi-programmed senescence and its pharmacologic inhibition. Cell Cycle 5: 2087-2102.

Blagosklonny MV. 2010. Revisiting the antagonistic pleiotropy theory of aging: TOR-driven program and quasi-program. Cell Cycle 9: 3151-3156.

Budovskaya YV, Wu K, Southworth LK, Jiang M, Tedesco P, Johnson TE, Kim SK. 2008. An elt-3/elt-5/elt-6 GATA transcription circuit guides aging in C. elegans. Cell 134: 291-303.

Chapin H, Okada M, Merz A, Miller D. 2015. Tissue-specific autophagy responses to aging and stress in C. elegans. Aging 7: 419-434.

Coburn C, Allman E, Mahanti P, Benedetto A, Cabreiro F, Pincus Z, Matthijssens F, Araiz C, Mandel A, Vlachos M et al. 2013. Anthranilate fluorescence marks a calcium-propagated necrotic wave that promotes organismal death in C. elegans. PLoS Biol 11: e1001613.

Cuervo A. 2004. Autophagy: in sickness and in health. Trends Cell Biol 14: 70-77.

de la Guardia Y, Gilliat AF, Hellberg J, Rennert P, Cabreiro F, Gems D. 2016. Run-on of germline apoptosis promotes gonad senescence in C. elegans. Oncotarget 7: 39082-39096.

Depina A, Iser W, Park S, Maudsley S, Wilson M, Wolkow C. 2011. Regulation of Caenorhabditis elegans vitellogenesis by DAF-2/IIS through separable transcriptional and posttranscriptional mechanisms. BMC Physiol 11: 11.

Depuydt G, Xie F, Petyuk VA, Shanmugam N, Smolders A, Dhondt I, Brewer HM, Camp DG, Smith RD, Braeckman BP. 2013. Reduced insulin/IGF-1 signaling and dietary restriction inhibit translation but preserve muscle mass in Caenorhabditis elegans. Mol Cell Proteomics 12: 3624-3639.

Dhondt I, Petyuk V, Cai H, Vandemeulebroucke L, Vierstraete A, Smith R, Depuydt G, Braeckman B. 2016. FOXO/DAF-16 activation slows down turnover of the majority of proteins in C. elegans. Cell Rep 16: 3028-3040.

Garigan D, Hsu A, Fraser A, Kamath R, Ahringer J, Kenyon C. 2002. Genetic analysis of tissue aging in *Caenorhabditis elegans:* a role for heat-shock factor and bacterial proliferation. Genetics 161: 1101-1112.

Ge L, Melville D, Zhang M, Schekman R. 2013. The ER-Golgi intermediate compartment is a key membrane source for the LC3 lipidation step of autophagosome biogenesis. eLife 2: e00947.

Gelino S, Hansen M. 2012. Autophagy - an emerging anti-aging mechanism. J Clin Exp Pathol Supp 4: 1-24.

Gems D. 2015. The aging-disease false dichotomy: understanding senescence as pathology. Front Genet 6: 212.

Gems D, de la Guardia Y. 2013. Alternative perspectives on aging in C. elegans: reactive oxygen species or hyperfunction? Antioxid Redox Signal 19: 321-329.

Gems D, Partridge L. 2013. Genetics of longevity in model organisms: Debates and paradigm shifts. Ann Rev Physiol 75: 621-644.

Grant B, Hirsh D. 1999. Receptor-mediated endocytosis in the Caenorhabditis elegans oocyte. Mol Biol Cell 10: 4311-4326.

Haithcock E, Dayani Y, Neufeld E, Zahand AJ, Feinstein N, Mattout A, Gruenbaum Y, Liu J. 2005. Age-related changes of nuclear architecture in Caenorhabditis elegans. Proc Natl Acad Sci USA 102: 16690-16695.

Harman D. 1956. Aging: A theory based on free radical and radiation chemistry. J Gerontol 11: 298-300.

Herndon L, Schmeissner P, Dudaronek J, Brown P, Listner K, Sakano Y, Paupard M, Hall D, Driscoll M. 2002. Stochastic and genetic factors influence tissue-specific decline in ageing *C. elegans*. Nature 419: 808-814.

Hopkinson J, Butte N, Ellis K, Smith O. 2000. Lactation delays postpartum bone mineral accretion and temporarily alters its regional distribution in women J Nutr 130: 777-783.

Huang C, Xiong C, Kornfeld K. 2004. Measurements of age-related changes of physiological processes that predict lifespan of *Caenorhabditis elegans*. Proc Natl Acad Sci USA 101: 8084-8089.

Huffman D, Barzilai N. 2010. Contribution of adipose tissue to health span and longevity. Interdiscip Top Gerontol 37: 1-19.

Hughes SE, Huang C, Kornfeld K. 2011. Identification of mutations that delay somatic or reproductive aging of Caenorhabditis elegans. Genetics 189: 341-356.

Jones K, Ashrafi K. 2009. Caenrhabditis elegans as an emerging model for studying the basic biology of obesity. Disease Models Mech 2: 224-229.

Kenyon C. 2010. The genetics of ageing. Nature 464: 504-512.

Kimble J, Sharrock, WJ. 1983. Tissue-specific synthesis of yolk proteins in *Caenorhabditis elegans*. Dev Biol 96: 189-196.

Kirkwood TBL, Rose MR. 1991. Evolution of senescence: late survival sacrificed for reproduction. Phil Trans R Soc London 332: 15-24.

Klapper M, Ehmke M, Palgunow D, Bohme M, Matthaus C, Bergner G, Dietzek B, Popp J, Doring F. 2011. Fluorescence-based fixative and vital staining of lipid droplets in Caenorhabditis elegans reveal fat stores using microscopy and flow cytometry approaches. J Lipid Res 52: 1281-1293.

Lapierre LR, Silvestrini MJ, Nunez L, Ames K, Wong S, Le TT, Hansen M, Melendez A. 2013. Autophagy genes are required for normal lipid levels in C. elegans. Autophagy 9: 278-286.

Leiser SF, Jafari G, Primitivo M, Sutphin GL, Dong J, Leonard A, Fletcher M, Kaeberlein M. 2016. Age-associated vulval integrity is an important marker of nematode healthspan. Age 38: 419-431.

Li Y, Na K, Lee HJ, Lee EY, Paik YK. 2011. Contribution of sams-1 and pmt-1 to lipid homoeostasis in adult Caenorhabditis elegans. J Biochem 149: 529-538.

Libina N, Berman J, Kenyon C. 2003. Tissue-specific activities of *C. elegans* DAF-16 in the regulation of lifespan. Cell 115: 489-502.

Luo S, Kleemann GA, Ashraf JM, Shaw WM, Murphy CT. 2010. TGF-beta and insulin signaling regulate reproductive aging via oocyte and germline quality maintenance. Cell 143: 299-312.

McGee MD, Day N, Graham J, Melov S. 2012. cep-1/p53-dependent dysplastic pathology of the aging C. elegans gonad. Aging 4: 256-269.

McGee MD, Weber D, Day N, Vitelli C, Crippen D, Herndon LA, Hall DH, Melov S. 2011. Loss of intestinal nuclei and intestinal integrity in aging C. elegans. Aging Cell 10: 699-710.

Murphy CT, McCarroll SA, Bargmann CI, Fraser A, Kamath RS, Ahringer J, Li H, Kenyon CJ. 2003. Genes that act downstream of DAF-16 to influence the lifespan of *C. elegans*. Nature 424: 277-284.

Na H, Zhang P, Chen Y, Zhu X, Liu Y, Liu Y, Xie K, Xu N, Yang F, Yu Y et al. 2015. Identification of lipid droplet structure-like/resident proteins in Caenorhabditis elegans. Biochim Biophys Acta 1853: 2481-2491.

Palikaras K, Mari M, Petanidou B, Pasparaki A, Filippidis G, Tavernarakis N. 2017. Ectopic fat deposition contributes to age-associated pathology in Caenorhabditis elegans. J Lipid Res 58: 72-80.

Podshivalova K, Kerr R, Kenyon C. 2017. How a mutation that slows aging can also disproportionately extend end-of-life decrepitude. Cell Reports 19: 441-450.

Riesen M, Feyst I, Rattanavirotkul N, Ezcurra M, Tullet J, Papatheodorou I, Ziehm M, Au C, Gilliat A, Hellberg J et al. 2014. MDL-1, a growth- and tumor-suppressor, slows aging and prevents germline hyperplasia and hypertrophy in C. elegans. Aging 6: 98-117.

Rodriguez JA, Marigorta UM, Hughes DA, Spataro N, Bosch E, Navarro A. 2017. Antagonistic pleiotropy and mutation accumulation influence human senescence and disease. Nat Ecol Evol 1: 55.

Rolls MM, Hall DH, Victor M, Stelzer EH, Rapoport TA. 2002. Targeting of rough endoplasmic reticulum membrane proteins and ribosomes in invertebrate neurons. Mol Biol Cell 13: 1778-1791.

Rompay L, Borghgraef C, Beets I, Caers J, Temmerman L. 2015. New genetic regulators question relevance of abundant yolk protein production in C. elegans. Sci Rep 5: 16381.

Seah NE, de Magalhaes Filho CD, Petrashen AP, Henderson HR, Laguer J, Gonzalez J, Dillin A, Hansen M, Lapierre LR. 2016. Autophagy-mediated longevity is modulated by lipoprotein biogenesis. Autophagy 12: 261-272.

Sharrock WJ, Sutherlin ME, Leske K, Cheng TK, Kim TY. 1990. Two distinct yolk lipoprotein complexes from Caenorhabditis elegans. J Biol Chem 265: 14422-14431.

Shi C, Runnels A, Murphy C. 2017. Mating and male pheromone kill Caenorhabditis males through distinct mechanisms. eLife 6: e23493.

Shintani T, Klionsky D. 2004. Autophagy in health and disease: a double-edged sword. Science 306: 990-995.

Shore D, Ruvkun G. 2013. A cytoprotective perspective on longevity regulation. Trends Cell Biol 23 409-420.

Steinbaugh M, Narasimhan S, Robida-Stubbs S, Moronetti Mazzeo L, Dreyfuss J, Hourihan J, Raghavan P, Operan T, Esmaillie R, Blackwell T. 2015. Lipid-mediated regulation of SKN-1/Nrf in response to germ cell absence. eLife 4: e07836.

Stout GJ, Stigter EC, Essers PB, Mulder KW, Kolkman A, Snijders DS, van den Broek NJ, Betist MC, Korswagen HC, Macinnes AW et al. 2013. Insulin/IGF-1-mediated longevity is marked by reduced protein metabolism. Mol Syst Biol 9: 679.

Thyagarajan B, Blaszczak A, Chandler K, Watts J, Johnson W, Graves B. 2010. ETS-4 is a transcriptional regulator of life span in Caenorhabditis elegans. PLoS Genet 6: e1001125.

Tian E, Wang F, Han J, Zhang H. 2009. epg-1 functions in autophagy-regulated processes and may encode a highly divergent Atg13 homolog in C. elegans. Autophagy 5: 608-615.

Tullet JM, Hertweck M, An JH, Baker J, Hwang JY, Liu S, Oliveira RP, Baumeister R, Blackwell TK. 2008. Direct inhibition of the longevity-promoting factor SKN-1 by insulin-like signaling in C. elegans. Cell 132: 1025-1038.

Van Raamsdonk JM, Hekimi S. 2010. Reactive oxygen species and aging in Caenorhabditis elegans: causal or casual relationship? Antioxid Redox Signal 13: 1911-1953.

Walther D, Kasturi P, Zheng M, Pinkert S, Vecchi G, Ciryam P, Morimoto R, Dobson C, Vendruscolo M, Mann M et al. 2015. Widespread proteome remodeling and aggregation in aging C. elegans. Cell 161: 919-932.

Wang H, Zhao Y, Ezcurra M, Gilliat A, Hellberg J, Benedetto A, Athigapanich T, Girstmair J, Telford M, Zhang Z et al. 2017. Monsters in the uterus: A parthenogenetic quasi-program causes teratoma-like tumors during aging in wild-type C. elegans. bioRxiv dx.doi.org/10.1101/174771.

Watts JL, Browse J. 2002. Genetic dissection of polyunsaturated fatty acid synthesis in Caenorhabditis elegans. Proc Natl Acad Sci USA 99: 5854-5859.

Wilhelm T, Byrne J, Medina R, Geisinger J, Hajduskova M, Tursun B, Richly H. 2017. Neuronal inhibition of the autophagy nucleation complex extends life span in post-reproductive C. elegans Genes and Development 31: 1561-1572.

Williams GC. 1957. Pleiotropy, natural selection and the evolution of senescence. Evolution 11: 398-411.

Zhao Y, Gilliat AF, Ziehm M, Turmaine M, Wang H, Ezcurra M. C. Y, Phillips G, McBay D, Zhang WB et al. 2017. Two forms of death in aging Caenorhabditis elegans. Nature Comm 8: 15458.

## Supplementary References

Chang J, Kumsta C, Hellman A, Adams L, Hansen M. 2017. Spatiotemporal regulation of autophagy during Caenorhabditis elegans aging. eLife 6: e18459.

Chapin H, Okada M, Merz A, Miller D. 2015. Tissue-specific autophagy responses to aging and stress in C. elegans. Aging 7: 419–434.

Dambroise E, Monnier L, Ruisheng L, Aguilaniu H, Joly J-S, Tricoire H, Rera M. 2016. Two phases of aging separated by the Smurf transition as a public path to death. Scientific Reports 6: 23523.

Depuydt G, Shanmugam N, Rasulova M, Dhondt I, Braeckman BP. 2016. Increased protein stability and decreased protein turnover in the C. elegans Ins/IGF-1 daf-2 mutant. J Gerontol 71: 1553–1559.

Depuydt G, Xie F, Petyuk VA, Shanmugam N, Smolders A, Dhondt I, Brewer HM, Camp DG, Smith RD, Braeckman BP. 2013. Reduced insulin/IGF-1 signaling and dietary restriction inhibit translation but preserve muscle mass in Caenorhabditis elegans. Mol Cell Proteomics 12: 3624–3639.

Dhondt I, Petyuk V, Bauer S, Brewer H, Smith R, Depuydt G, Braeckman B. 2017. Changes of protein turnover in aging Caenorhabditis elegans. Mol Cell Proteomics 16: 1621–1633.

Dhondt I, Petyuk V, Cai H, Vandemeulebroucke L, Vierstraete A, Smith R, Depuydt G, Braeckman B. 2016. FOXO/DAF-16 activation slows down turnover of the majority of proteins in C. elegans. Cell Rep 16: 3028–3040.

Gelino S, Chang J, Kumsta C, She X, Davis A, Nguyen C, Panowski S, Hansen M. 2016. Intestinal autophagy improves healthspan and longevity in C. elegans during dietary restriction. PLoS Genet 12: e1006135.

Gelino S, Hansen M. 2012. Autophagy - an emerging anti-aging mechanism. J Clin Exp Pathol Supp 4: 1–24.

Hansen M, Chandra A, Mitic LL, Onken B, Driscoll M, Kenyon C. 2008. A role for autophagy in the extension of lifespan by dietary restriction in C. elegans. PLoS Genet 4: e24.

Hashimoto Y, Ookuma S, Nishida E. 2009. Lifespan extension by suppression of autophagy genes in Caenorhabditis elegans. Genes Cells 14: 717–726.

Jia K, Thomas C, Akbar M, Sun Q, Adams-Huet B, Gilpin C, Levine B. 2009. Autophagy genes protect against Salmonella typhimurium infection and mediate insulin signaling-regulated pathogen resistance. Proc Natl Acad Sci SA 106: 14564–14569.

Kang C, Avery L. 2008. To be or not to be, the level of autophagy is the question: Dual roles of autophagy in the survival response to starvation. Autophagy 4: 82–84.

Kimble J, Sharrock, WJ. 1983. Tissue-specific synthesis of yolk proteins i. Caenorhabditis elegans. Dev Biol 96: 189–196.

Kumsta C, Chang J, Schmalz J, Hansen M. 2017. Hormetic heat stress and HSF-1 induce autophagy to improve survival and proteostasis in C. elegans. Nat Commun 8: 14337.

Lapierre L, Gelino S, Meléndez A, Hansen M. 2011. Autophagy and lipid metabolism coordinately modulate life span in germline-less C. elegans. Curr Biol 21: 1507–1514.

Meléndez A, Tallóczy Z, Seaman M, Eskelinen E-L, Hall DH, Levine B. 2003. Autophagy genes are essential for dauer development and life-span extension i. C. elegans. Science 301: 1387–1391.

Narita M, Young A, Arakawa S, Samarajiwa S, Nakashima T, Yoshida S, Hong S, Berry L, Reichelt S, Ferreira M et al. 2011. Spatial coupling of mTOR and autophagy augments secretory phenotypes. Science 332: 966–970.

Shintani T, Klionsky D. 2004. Autophagy in health and disease: a double-edged sword. Science 306: 990–995.

Stout GJ, Stigter EC, Essers PB, Mulder KW, Kolkman A, Snijders DS, van de. Broek NJ, Betist MC, Korswagen HC, Macinnes A. et al. 2013. Insulin/IGF-1-mediated longevity is marked by reduced protein metabolism. Mol Syst Biol 9: 679.

Visscher M, De Henau S, Wildschut M, van Es R, Dhondt I, Michels H, Kemmeren P, Nollen E, Braeckman B, Burgering B et al. 2016. Proteome-wide changes in protein turnover rates in C. elegans models of longevity and age-related disease. Cell Rep 16: 3041–3051.

Wilhelm T, Byrne J, Medina R, Geisinger J, Hajduskova M, Tursun B, Richly H. 2017. Neuronal inhibition of the autophagy nucleation complex extends life span in post-reproductive C. elegans Genes Develop 31: 1561–1572.

Zhang H, Chang J, Guo B, Hansen M, Jia K, Kovács A, Kumsta C, Lapierre L, Legouis R, Lin L et al. 2015. Guidelines for monitoring autophagy in Caenorhabditis elegans. Autophagy 11: 9–27.

